# Nutrient levels control root growth responses to high ambient temperature in plants

**DOI:** 10.1101/2023.08.04.552051

**Authors:** Sanghwa Lee, Julia Showalter, Ling Zhang, Gaëlle Cassin-Ross, Hatem Rouached, Wolfgang Busch

## Abstract

Global warming will lead to significantly increased temperatures on earth. Plants respond to high ambient temperature with altered developmental and growth programs, termed thermomorphogenesis. Here we show that thermomorphogenesis is conserved in Arabidopsis, soybean, and rice and that it is linked to a decrease in the levels of the two macronutrients nitrogen and phosphorus. We also find that low external levels of these nutrients abolish root growth responses to high ambient temperature. We show that in Arabidopsis, this is due to the function of the transcription factor *ELONGATED HYPOCOTYL 5* (*HY5*) and its transcriptional regulation of the transceptor *NITRATE TRANSPORTER 1.1* (*NRT1.1*). Soybean and Rice homologs of these genes are expressed consistently with a conserved role in regulating temperature responses in a nitrogen and phosphorus level dependent manner. Overall, our data show that root thermomorphogenesis is a conserved feature in species of the two major groups of angiosperms, monocots and dicots, that it leads to a reduction of nutrient levels in the plant, and that it is dependent on sufficient environmental nutrient supply, a regulatory process mediated by the HY5-NRT1.1 module.

## Introduction

The recent rise in global temperature is largely due to human activities and is predicted to continue. Most optimistic scenarios predict a 1.5°C increase of global average temperatures by mid-century, while the middle of the road scenarios predict 2.7°C increase by the end of the century (IPCC, 2022). Temperature profoundly affects biological systems due to its effect on the free energy for biochemical reactions according to the basic principles of thermodynamics (Held and Sadowski, 2016; Ibanez et al., 2017). Plants don’t regulate their internal temperature and due to their sessile nature, they are very sensitive to climate change (Schleuning et al., 2016). Plants respond to high ambient temperature with a developmental program termed thermomorphogenesis. The hall-mark phenotypes of thermomorphogenesis are elongated tissues including hypocotyl, petiole and root, hyponastic growth, stomatal development, and early flowering (Gaillochet et al., 2020; Lee et al., 2021a; Martins et al., 2017; Vu et al., 2019).

There are two transcription factors serving as central hub in thermomorphogenesis, PHYTOCHROME INTERACTING FACTOR 4 (PIF4) and ELONGATED HYPOCOTYL 5 (HY5) both of which were originally identified as light signaling components (Huq and Quail, 2002; Koini et al., 2009; Lee *et al*., 2021a; Oyama et al., 1997). PIF4 has a major role in the thermomorphogenesis of the shoot, which also involves other PIFs such as PIF1, 3, 5, and 7 (Burko et al., 2022; Fiorucci et al., 2019; Koini *et al*., 2009; Kumar et al., 2012; Lau et al., 2018; Lee et al., 2021b). HY5 plays a major role in root thermomorphogenesis, which regulates primary root length at the early seedling stage (Gaillochet *et al*., 2020; Lee *et al*., 2021a).

Another key factor for growth and development is nutrient availability. There is a strong interaction of nutrient availability and temperature for determining growth (G-Yull Rhee, 1981). Conversely, nutrient content is affected by elevated temperature and increased CO_2_ levels (Bouain et al., 2022; Carvalho et al., 2020; Cassan et al., 2023). Nitrogen (N), Phosphorus (P), and Potassium (K) are three macronutrients which are commonly used for agricultural fertilizer and enhance growth. Due to the importance of macronutrient uptake, nutrient signaling has been widely studied. One of the most well-studied genes in nitrogen uptake is *NITRATE TRANSPORTER 1.1* (*NRT1.1*), which encodes for a dual-affinity nitrate transporter and nitrate sensor (Krouk et al., 2010; Tsay et al., 1993; Ye et al., 2019). Furthermore, *OsNRT1.1B*, which is a functional homolog of *AtNRT1.1* in rice, has been showed to integrate N and P signaling (Hu et al., 2019), suggesting that the role of NRT1.1 as a master regulator of N and P might be conserved across the plant kingdom.

Here, we show that shoot and root thermomorphogenesis are conserved among Arabidopsis (*Arabidopsis thaliana*), soybean (*Glycine max*), and rice (*Oryza sativa*). We find that this is linked to decreased N and P levels in plant tissues at higher temperature. Conversely, low levels of N and P in the medium abolished thermomorphogenesis in Arabidopsis. We found this to be regulated by a module constituted by the thermomorphogenesis key regulator *HY5* and the nitrogen transceptor *NRT1.1*.

## Result

### Plants show conserved shoot and root thermomorphogenesis that goes along with decreased N and P levels in plant tissue

Root thermomorphogenesis studies have been largely restricted to Arabidopsis. We therefore wanted to compare this to the response in other species. For this, we grew Arabidopsis, rice, and soybean seedlings at ambient and elevated temperatures. For Arabidopsis, Col-0 wild type seedlings were grown at either 21LJ or 28LJ for 5 days after 4 days of germination at 21LJ (Fig. 1a, b). Soybean (Williams 82 variety) and rice (Kitaake ecotype) were grown at either 28LJ or 33LJ for 1 or 2 weeks for soybean and rice respectively after 7 days of germination at 28LJ (Fig. 1c-f). Similar to the reported Arabidopsis shoot and root thermomorphogenesis (Fig.1 a, b; (Koini *et al*., 2009; Lee *et al*., 2021a)), both soybean and rice showed longer shoots and roots at higher temperature, indicating that plants have a conserved elongation mechanism at higher temperature.

**Figure 1.**
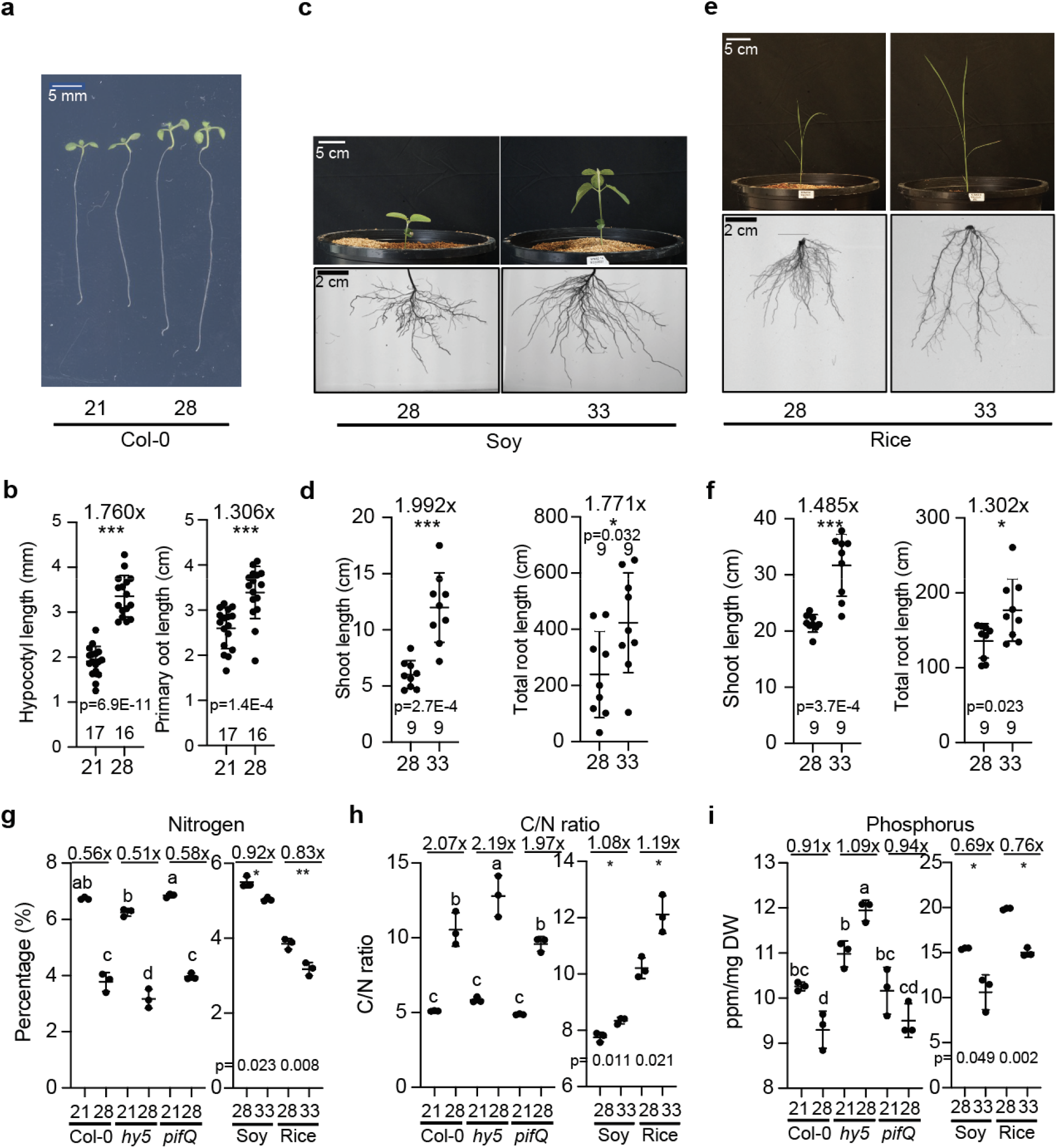
Plant developmental responses to high ambient temperatures are conserved and are linked to altered nutrient levels. **a-f** Phenotypes of Arabidopsis (Col-0; **a, b**), soybean (Williams82; **c, d**), and rice (Kitaake; **e, f**) at normal and higher temperatures. Arabidopsis seedlings were grown for 4 days on 1/2ms plates at 21LJ and then kept either at 21LJ or 28 LJ for 5 additional days. Rice and soybean seedlings were grown for 1 week at 28 LJ and then either kept at 28 LJ or 33 LJ for additional 2 weeks for rice and 1 week for soybean respectively. Scatter dot plot shows average difference of primary root length for Arabidopsis, and total root length for soybean and rice, and the number of plants. P-value from one-sided Student’s t-test. **g, h** Nitrogen (**g**) and C/N ratio (**h**) in Arabidopsis shoots (Col-0, *hy5-215*, and *pifQ*), soybean shoots, and rice shoots using CN analysis. **i** Phosphorus in Arabidopsis shoots, Soybean shoots, and rice shoots using MP-AES. Letters a-d indicate statistically significant differences based on one-way ANOVA analysis with Tukey’s HSD test. Asterisks indicate statistically significant difference using Student’s t-test; *p<0.05, **p<0.01, and ***p<0.001. P-value from one-sided Student’s t-test. Average fold difference of each group is indicated in the top region of the plot. Shoot parts from 4-week-old plants from Arabidopsis, soybean, and rice plants were used for the nutrient analyses. Plots indicate mean (horizontal line) and standard deviation (error bars).

Previous studies had exposed temperature dependent gene expression changes of gene clusters related to nitrogen and organic acid related processes (Supplementary Table 1; (Gaillochet *et al*., 2020; Lee *et al*., 2021a). Furthermore, HY5 has been found to regulate nitrogen and iron signaling (Guo et al., 2021; Huang et al., 2015). We therefore hypothesized that nutrient composition or uptake could be changed at higher temperature. To test this hypothesis, we analyzed the nutrient composition of shoots of 4 week-grown plants of Arabidopsis, rice, and soybean (Fig. 1g-i, Supplementary Fig. 1). We also included Arabidopsis *hy5-215* and *pifQ* mutants in this analysis. Interestingly, levels of N and P were decreased in Arabidopsis, soybean, and rice at higher temperature (Fig. 1g-i), while other nutrients showed less consistent pattern across the species at higher temperature (Supplementary Fig. 1). Interestingly, the *hy5-215* mutant plants showed different patterns of N and P level changes, whereas *pifQ* was similar to Col-0 at high ambient temperature, indicating that *HY5* might be involved in the alteration of N and P levels at high ambient temperature (Fig. 1g-i). Overall, this suggested that the levels of N and P in plants are regulated in response to elevated temperature, and that HY5 might be involved in this regulation.

### HY5 integrates temperature and N-P signaling

Since we found *HY5* to be involved in altering plant nutrient levels at high ambient temperature, we searched for a target downstream of *HY5* that could explain its function. HY5 is a bZIP protein transcription factor, which binds to several DNA sequence motifs including G-box (CACGTG) and CACGT motifs (Burko et al., 2020; Lee et al., 2007). Published ChIP-seq data showed that HY5 binds to the promoter region of genes that are involved in nutrient related responses such as those that we had identified using our RNAseq to be related to nitrogen and organic acids (Burko *et al*., 2020; Lee *et al*., 2007; Lee *et al*., 2021a); Supplementary Table. 1). We identified genes that were bound by HY5 according to the ChIP-seq data (Burko *et al*., 2020) and that are in the N-P signaling pathway. These genes included *NIGT1.1, HHO2, NLP7, NRT1.1, NRT1.5*, *NRT2.1*, *NIA1*, and *LBD37* (Alvarez et al., 2020; Chen et al., 2012; Krouk *et al*., 2010; Remans et al., 2006; Wang et al., 2020; Yu et al., 1998). To examine whether HY5 directly binds to the promoter of these genes in our growth conditions (the published ChIP-seq data were obtained under different light conditions), we performed Chromatin Immunoprecipitation qPCR (ChIP-qPCR) using 4 days 21LJ grown *pHY5:HY5-GFP* seedlings with additional 5 days treated in either 21LJ or 28LJ (Fig. 2a, b, supplementary Fig. 2a, b). Interestingly, not all but only a few target promoters such as *NIGT1.1, HHO2, NLP7,* and *NRT1.1* were enriched at high ambient temperature under our growth conditions, indicating that HY5 directly binds to N-P signaling genes and regulates expression in a temperature dependent manner. To test whether the transcription levels of those genes are altered at high ambient temperature, we performed qPCR of root and shoot tissues (Fig. 2c, supplementary Fig. 2c). Consistent with our ChIP-qPCR data, temperature dependent HY5 enriched target genes such as *NIGT1.1, HHO2, NLP7,* and *NRT1.1* were significantly downregulated while the genes for which we hadn’t found ChIP-qPCR enrichment, such as *NRT1.5*, *NRT2.1, NIA1, and LBD37* were not altered at high ambient temperatures. Among the genes that were bound by *HY5* and downregulated, *NRT1.1* stood out as its transcript levels were downregulated at high ambient temperature in the root of Col-0, but didn’t change its expression level in the roots of *hy5-215* mutant plants. This suggests that the transcript level of *NRT1.1* is tightly regulated by HY5 in the root in a temperature dependent manner. While the transcript level of *NIA1* was altered in *hy5-215* mutant, there was no indication of a change in HY5 binding at high ambient temperature according to our ChIP-qPCR (Supplementary Fig. 2a-c), suggesting that *NIA1* transcript level is altered indirectly. Furthermore, the transcript level of *NRT1.5* decreased at high ambient temperature both in Col-0 and *hy5-215* mutant (Supplementary Fig. 2a-c), indicating that another component might be responsible for regulating *NRT1.5* transcript levels at high ambient temperature. Overall, the data suggest that HY5 regulates root thermomophogenesis and nutrient composition by repressing the N-P signaling genes and by directly regulating key genes such as *NRT1.1*.

**Figure 2.**
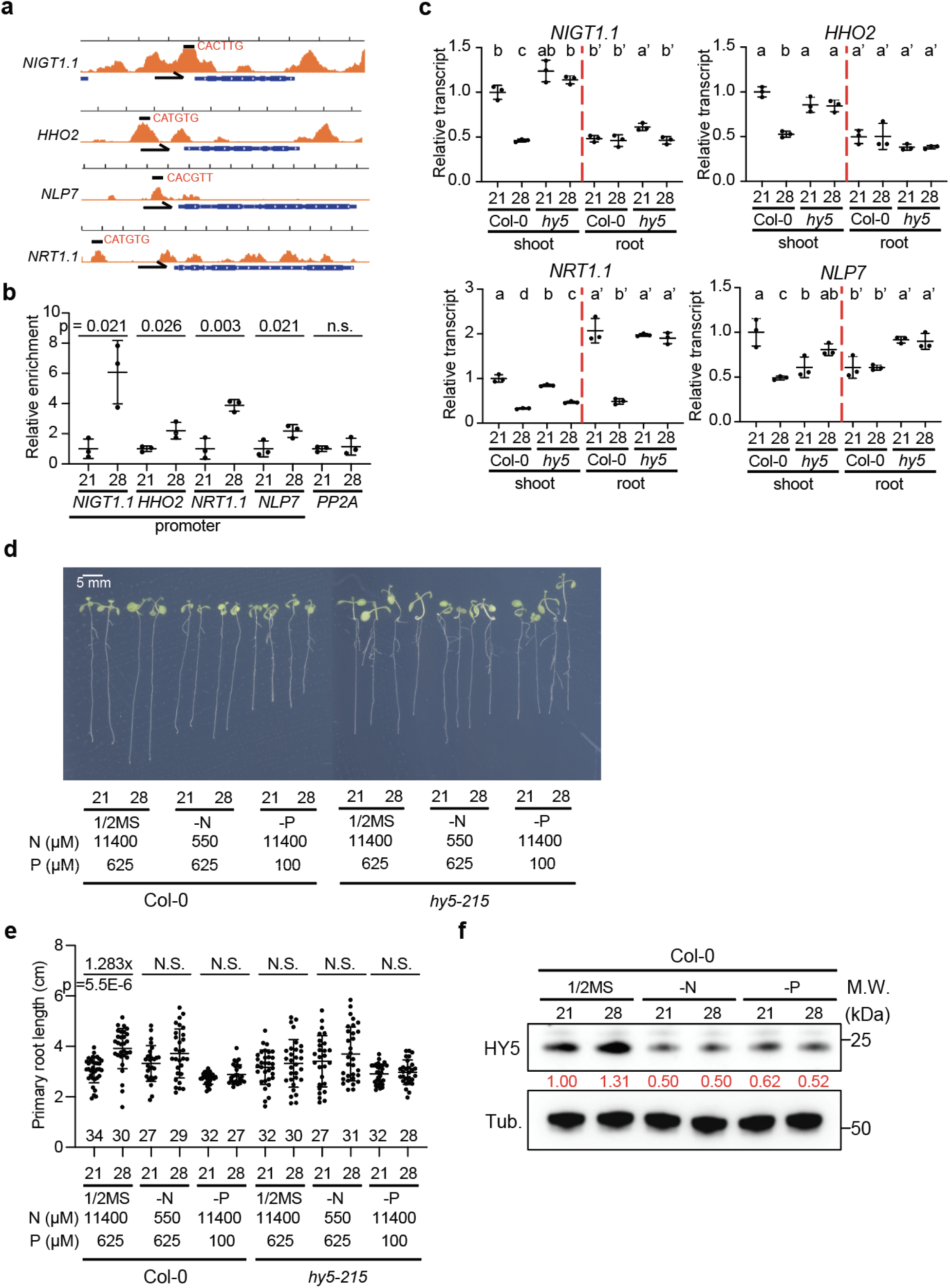
HY5 directly regulates N-P signaling genes at high ambient temperature. **a** IGV image of HY5 ChIP-seq data from Burko et al., (2020) of selected N-P signaling genes with transcription direction and binding motif. **b** Scatter dot plot of ChIP-qPCR results at normal and high ambient temperature of promoter regions of five different genes. **c** Scatter dot plot of qPCR results at normal and high ambient temperature of four different genes using Col-0 and *hy5-215* shoot and root samples of seedlings. Relative transcript level was normalized using *PP2A* as a control and to the expression levels in the shoot . **d, e** Phenotypes of Col-0 and *hy5-215* grown on different media at high ambient temperature. Letters a-d indicate statistically significant differences based on one-way ANOVA analysis with Tukey’s HSD test. Shoot and root samples were analyzed separately. P value from one-sided Student’s t-test. Average fold difference of each group is indicated in the top region of the plot. **f** Western blot analysis using native HY5 antibody. Red number indicates the relative signal intensity divided by HY5 signal to Tubulin. Scatter dot plots indicate mean (horizontal line) and standard deviation (error bars).

Because *HY5* was involved in determining N-P levels at high ambient temperature (Fig. 1g-i), we hypothesized that *HY5* levels might be affected by N-P deficient conditions at high ambient temperature. To test this hypothesis, Col-0 and *hy5-215* plants were grown for 5 days at either 21LJ or 28LJ after 4 days for germination in 21LJ in three different media conditions: 1/2MS (N:11400 μM, P: 625 μM), mildly nitrogen deficient (N: 550 μM), and mildly phosphorus deficient (P: 100 μM) (Gruber et al., 2013). We then analyzed protein abundance of HY5 using Western Blots of root samples (Fig. 2f). Similar to a short time of exposure to high ambient temperature for 4 hours, leading to HY5 accumulation (Lee *et al*., 2021a), HY5 protein level was accumulated even after long time of temperature exposure for 5 days in nutrient sufficient ½ MS media, however, HY5 protein levels did not change in N or P deficient media. We then tested whether the altered HY5 level affected with altered root thermomorphogenesis in these three different conditions (Fig. 2d, e). Root thermomorphogenesis was observed only in ½ MS grown seedlings, which display higher HY5 protein levels at high ambient temperature (Fig. 2c). Grown on N or P deficient media, seedlings did not show longer primary root at high ambient temperature. This was similar to the response of the *hy5-215* mutant, in which there is no HY5 protein produced (Fig. 2d, e). Overall, these data suggested that HY5 is essential for temperature mediated primary root elongation and that the lack of temperature dependent root growth response in N or P deficient medium is due to the lack of increased HY5 levels.

We then tested the effect of excessive amount of N or P on root thermomorphogenesis. For this, Col-0 seedlings were grown in N or P excessive media at high ambient temperature (Supplementary Fig. 3). Col-0 seedlings did not have exaggerated root thermomorphogenesis in N or P excessive media at high ambient temperature when comparing it to 1/2MS conditions, indicating that excessive amounts of N or P do not affect root elongation at high ambient temperature. Taken together, these data suggest that sufficient amounts of N and P is necessary for HY5 mediated root thermomorphogenesis.

### NRT1.1 integrates N and P dependent root thermomorphogenesis

NRT1.1 has been shown to be a key component in nitrate signaling and is regulated by nitrate and phosphate level, as well as by auxin in root development (Fang et al., 2021; Krouk *et al*., 2010; Medici et al., 2015). As *NRT1.1* transcript levels were decreased at high ambient temperature and were tightly regulated by HY5 in a root specific manner (Fig. 2c), we performed qPCR using roots of seedlings grown on nutrient sufficient or deficient media (Fig. 3a). These plants were grown at either 21LJ or 28LJ for 5 days after they had germinated and grown for 4 days at 21LJ. Seedlings grown on nutrient sufficient medium displayed significantly decreased transcript levels of *NRT1.1* at high ambient temperatures. Seedlings grown on N or P deficient media showed a decreased level of NRT1.1 transcript at 21°C (albeit not as low as those grown on at 28°C and nutrient sufficient medium) and this was not further decreased in high ambient temperature. To investigate these transcriptional changes in higher detail, we analyzed seedlings of the *pNRT1.1:GFP* reporter line (Guo et al., 2001) grown at 21°C or 28°C, to assess tissue specific expression changes in different nutrient conditions using confocal microscopy (Fig. 3b, c). Reflecting the data from the qPCR, the *pNRT1.1:GFP* signal was decreased in the root apex at high ambient temperature in roots grown on nutrient sufficient medium. The GFP signal was much lower in roots grown on N or P deficient media at both temperatures (Fig. 3b, c). Overall, these data suggest that sufficient amounts of N and P are necessary for the suppression of *NRT1.1* transcript level at high ambient temperature.

**Figure 3.**
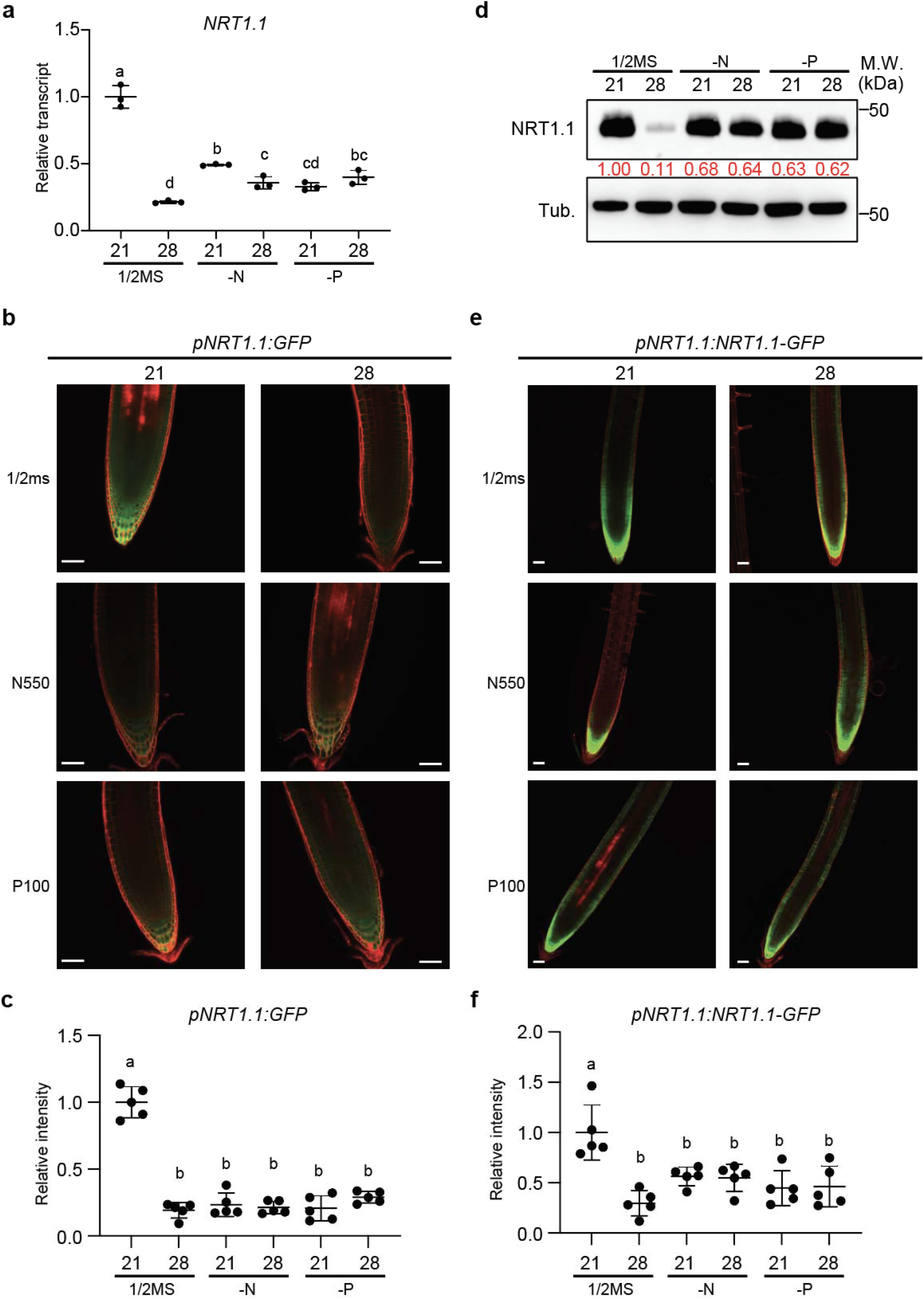
NRT1.1 integrates N-P signaling and root thermomorphogenesis. **a** Scatter dot plot of qPCR results at normal and high ambient temperature grown Col-0 seedlings grown on different media (1/2MS, -N, -P). Only root samples were used for the analysis. The relative transcript level of *NRT1.1* was normalized by the expression levels of *PP2A* and to the expression levels in the shoot. **b** Confocal microscopy images of *pNRT1.1:GFP* transgenic line at normal and high ambient temperature grown on different media (1/2MS, -N, -P). **c** Scatter dot plot of the signals quantified from confocal microscopy images of **b**. Quantification of the signal intensity was measured. **d** Western blot analysis of Col-0 seedling roots using native NRT1.1 antibody. Red number indicates the relative signal intensity divided by NRT1.1 signal to Tubulin. **e** Confocal microscopy images of *pNRT1.1:NRT1.1-GFP* transgenic line at normal and high ambient temperature grown on different media (1/2ms, -N, -P). Scale bar indicates 50 μm. **f** Scatter dot plot of the signals quantified from confocal microscopy images of **e**. Letters a-d indicate statistically significant differences based on one-way ANOVA analysis with Tukey’s HSD test. Media concentrations includes: 1/2MS (N:11400 μM, P: 625 μM), mildly nitrogen deficient (N: 550 μM), and mildly phosphorus deficient (P: 100 μM). Scatter dot plots indicate mean (horizontal line) and standard deviation (error bars).

As *NRT1.1* transcript level is altered in different media and temperature conditions (Fig. 3a-c), we measured total root protein levels of NRT1.1 using a native NRT1.1 antibody (Fig. 3d). Similar to the transcriptional response, NRT1.1 protein levels decreased at high ambient temperature under nutrient sufficient conditions (Fig. 2c, Fig. 3a). This indicates that both transcription and protein level of NRT1.1 are decreased at high ambient temperature. In -N and -P conditions, protein levels of NRT1.1 were not strikingly different from nutrient sufficient conditions, indicating that the observed differences in transcription levels were not reflecting protein levels. Importantly, the decrease of NRT1.1 protein level observed in nutrient sufficient medium at high ambient temperature, was abolished in -N and -P media. To assess this regulation at the protein level at cellular resolution, we utilized *pNRT1.1:NRT1.1-GFP* transgenic plants (Fig. 3e, f (Krouk *et al*., 2010)). NRT1.1 protein level was decreased in the root apex in ½ MS media at high ambient temperature, while N or P deficient media grown seedlings did not show decreased *NRT1.1-GFP* signal in the root tip (Fig. 3e, f). Overall, the data suggest that changes in NRT1.1 transcript and protein level are necessary for root thermomorphogenesis.

### The *HY5*-*NRT1.1* regulatory module is required for the interaction of root thermomorphogenesis and N-P levels

As *NRT1.1* transcript level was tightly regulated by HY5 in the root at high ambient temperatures (Fig. 2c), we examined if there is a *HY5* feedback loop from NRT1.1 on *HY5*. qPCR and Western blot analyses (Fig. 4a, b) showed that *HY5* transcript level showed similar patterns in the *NRT1.1* loss of function mutant line (*chl1-5)* compared to Col-0, where *HY5* transcript level is induced at high ambient temperature in a root specific manner. This indicated that there is no significant feedback from NRT1.1 on the *HY5* transcript level. To test this at the protein level we performed western blots (Fig. 4b). Interestingly, HY5 protein level was more accumulated in *chl1-5* mutant roots in all our conditions, suggesting that *NRT1.1* regulates HY5 protein stability via a direct or indirect unknown mechanism. Furthermore, NRT1.1 protein accumulated to a higher extent in *hy5-215* mutant roots at high ambient temperature, similarly to the transcript level, indicating that HY5 has a major role to suppress *NRT1.1* transcript level and thus regulates NRT1.1 protein level at high ambient temperature.

**Figure 4.**
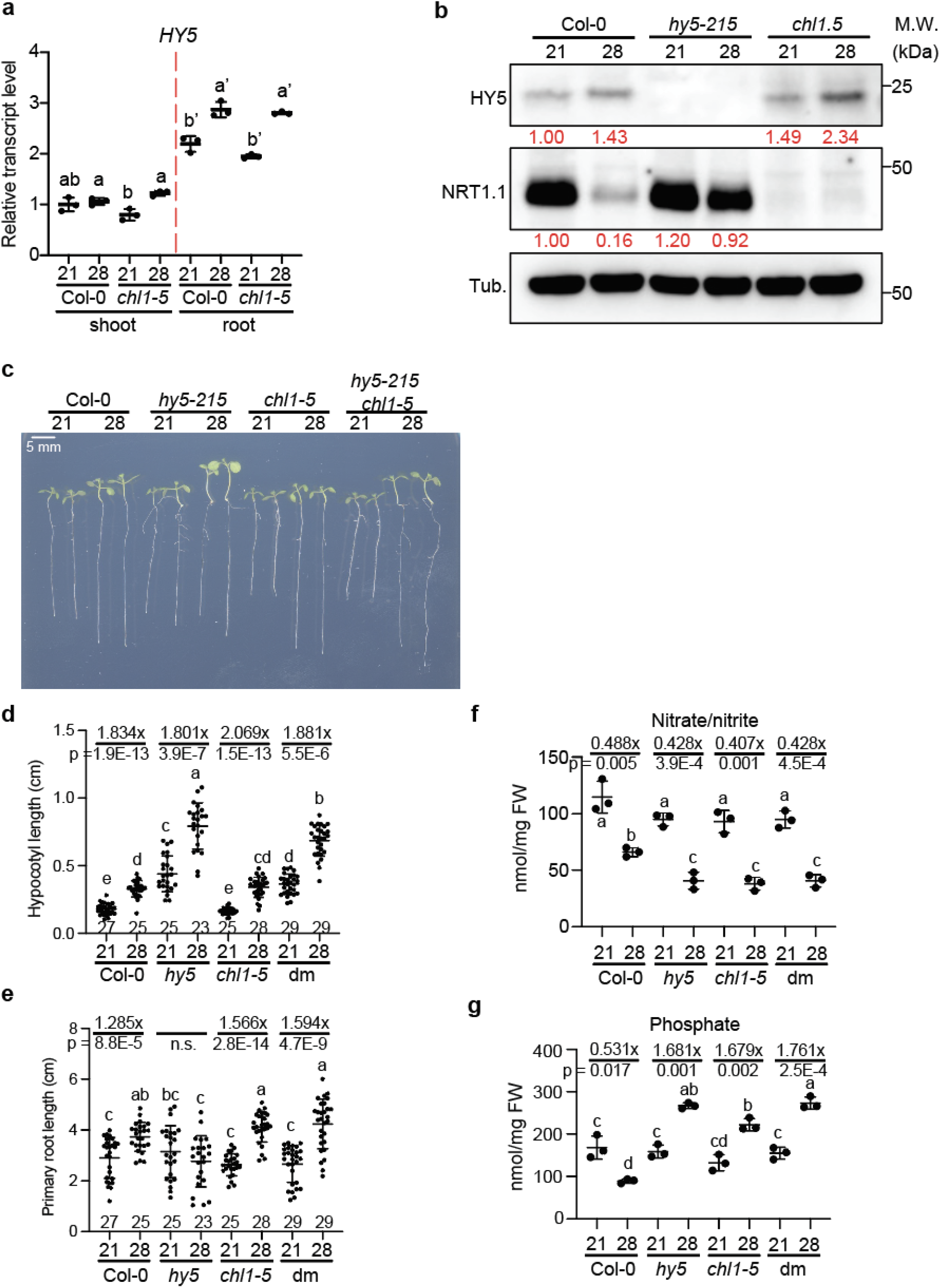
*HY5* and *NRT1.1* regulate root thermomorphogenesis and N-P alteration at high ambient temperature. **a-e** Phenotypic analyses of Col-0, *hy5-215*, *chl1-5*, *hy5-215 chl1-5* double mutant at high ambient temperature. **b-e** Scatter dot plot of phenotypic analyses, hypocotyl length (**b**), root length (**c**), nitrate and nitrite composition (**d**), and phosphate composition (**e**). For nitrate/nitrite and phosphate composition analyses, seeds of each genotype were grown on soil for 2 weeks at 21_J and transferred into either 21_J or 28_J for additional 5 weeks. Then the leaves were used for the analyses. **f** qPCR results of *HY5* transcript level at normal and high ambient temperature using Col-0 and *chl1-5* seedlings with separated samples of shoot and root. Relative transcript level was normalized using *PP2A* as a control and to the expression levels in the shoot. **g** Western blot analyses of NRT1.1 and HY5 using Col-0, *hy5-215*, and *chl1-5*. Red number indicates the relative signal intensity divided by NRT1.1 or HY5 signal to Tubulin. Letters a-d indicate statistically significant differences based on one-way ANOVA analysis with Tukey’s HSD test. Shoot and root samples were analyzed separately. P value from one-sided Student’s t-test. Average fold difference of each group is indicated in the top region of the plot. Scatter dot plots indicate mean (horizontal line) and standard deviation (error bars).

The direct binding of HY5 to the *NRT1.1* promoter and the transcriptional regulation of *NRT1.1* expression level by *HY5* (Fig. 2a-c) indicated that HY5 is upstream of NRT1.1. To genetically test this, we obtained mutant lines of *HY5* (*hy5-215)* and *NRT1.1* (*chl1-5)*, as well as the double mutant (*hy5-215 chl1-5*). We then assessed hypocotyl and primary root length, as well as N-P composition at standard and high ambient temperatures (Fig. 4c-g). As reported previously (Gaillochet *et al*., 2020; Lee *et al*., 2021a), *hy5-215* showed a thermo-insensitive root phenotype. The *chl1-5* mutant showed an increased root elongation phenotype at high ambient temperature compared to that of Col-0, while having the similar pattern of shoot thermomorphogenesis (Fig. 4c, d). Consistent with our data that HY5 represses *NRT1.1* expression in a root specific manner at high ambient temperature (Fig. 2c), *hy5-215 chl1-5* double mutant plants showed similar phenotypes to *hy5-215* in the shoot and *chl1.5* in the root respectively (Fig. 4c-e). This shows that *chl1-5* is epistatic to *hy5-215* in root thermomorphogenesis. To assess whether the root thermomorphogenesis phenotypes were linked to N-P level alterations at high ambient temperature, we measured nitrate/nitrite and phosphate levels in the mutants (Fig. 4f, g). Consistent with the MP-AES results (Fig. 1g, i), Col-0 and *hy5-215* showed different patterns of N-P level alteration at high ambient temperature. Interestingly, *chl1-5* and *hy5-215 chl1-5* double mutant plants showed similar N-P pattern with *hy5-215*, which indicates that the *NRT1.1* alteration upon temperature is critical for N-P level alteration at high ambient temperature. Taken together, these data suggest that *HY5-NRT1.1* regulatory mechanism has an important role to regulate root thermomorphogenesis and N-P level alteration at high ambient temperature.

### The *HY5*-*NRT1.1* regulatory mechanism alters global gene expression at high ambient temperature

Because the transcript level of *NRT1.1* was down-regulated by HY5 in a root specific manner at high ambient temperature, we performed RNAseq to examine the global gene expression pattern at high ambient temperature using Col-0, *hy5-215*, and *chl1-5* plants. These plants were grown at either 21LJ or 28LJ for additional 5 days after 4 days of germination at 21LJ. Consistent with our qPCR results, *NRT1.1* was among the most downregulated gene in Col-0 roots when comparing 28LJ to 21LJ (Fig. 5a).

**Figure 5.**
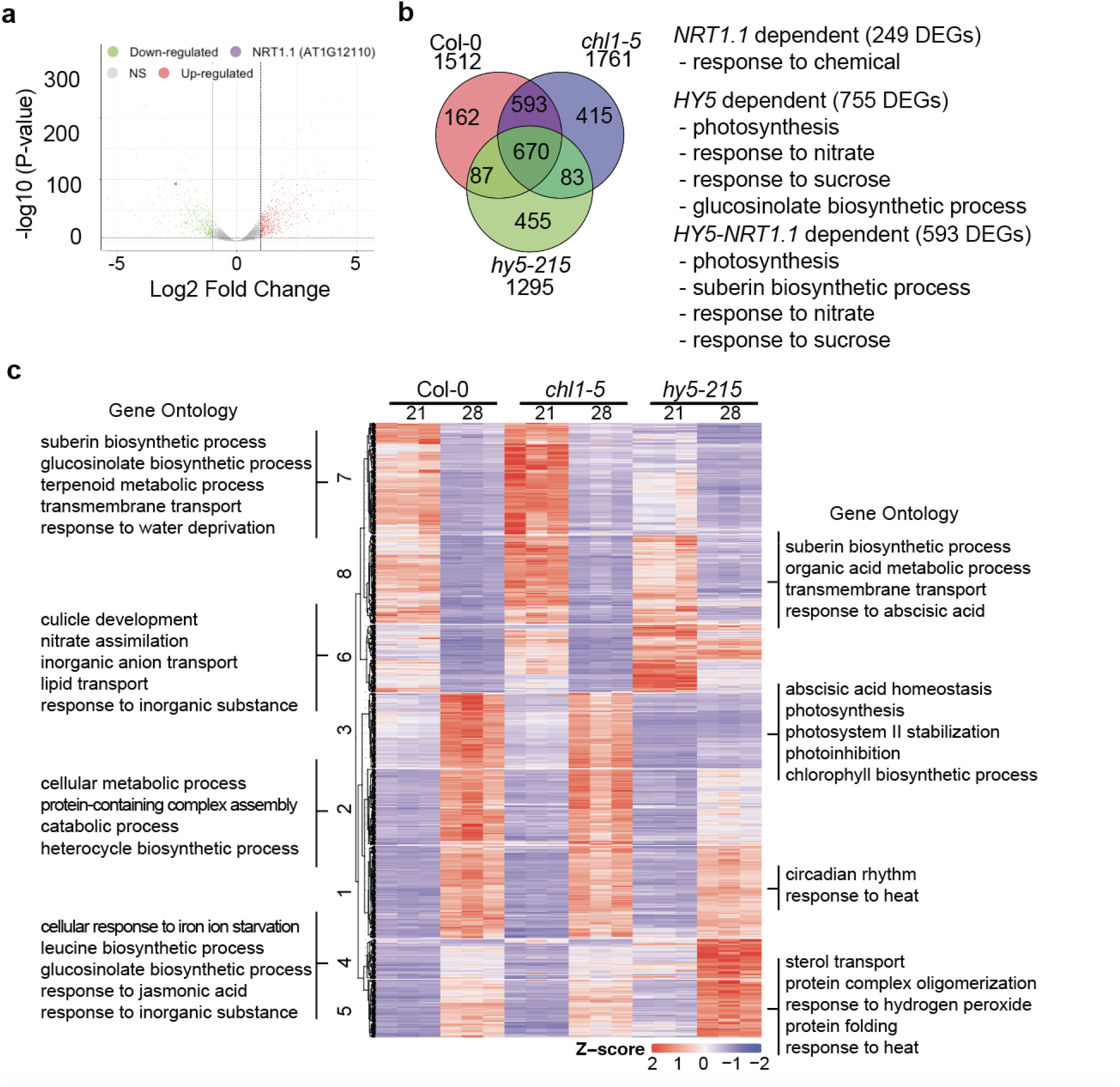
*HY5-NRT1.1* regulatory mechanism controls global gene expression at high ambient temperature. **a** Volcano plot of RNAseq using Col-0 root samples. Col-0 seedlings were grown for 4 days in 21LJ and then transferred either 21LJ or 28LJ for additional 5 days and collected. Threshold of P value is 0.05, and Log2 Fold change threshold is −1 and 1. *NRT1.1* is labeled with purple dot. **b** Venn diagram of root Differentially Expressed Genes (DEGs) in Col-0, *chl1-5,* and *hy5-215* mutant at high ambient temperature. Gene Ontology (GO) analysis clusters within each group are described next to the Venn diagram. **c** Heatmap analysis shows that Col-0, *chl1-5,* and *hy5-215* mutant have different expression pattern. Representative GO analyses of each cluster are noted. Three biological repeats were performed for RNAseq analysis.

Many genes were differentially expressed in these three genotypes in response to temperature: 1512, 1761, and 1295 DEGs in Col-0, *chl1-5*, and *hy5-215* respectively (Fig. 5b). More than 83% of root Col-0 DEGs were shared between Col-0 and *chl1-5*, indicating that despite the hypersensitive root thermomorphogenesis *chl1-5*, the mutant mounts a very similar response. When conducting a Gene Ontology (GO) analysis of the 249 *NRT1.1* dependent genes (DEGs in Col-0 but not in *chl1-5*), the significant GO-term “response to chemical” was significantly enriched. Also, almost half of the root Col-0 DEGs were *HY5* dependent (755 genes out of 1512 DEGs, >49.9%). This set of genes was enriched for the GO categories related to response to nutrients such as nitrate and sucrose. We reasoned that set of thermomorphogenesis genes would be genes that were not differentially expressed in *hy5-215* mutants (as there is no root growth response to temperature) but still occurs in *chl1-5* (this mutant as well as the *hy5-215/chl1-5* double mutant still shows a root growth response to elevated temperature). This was a set of 593 genes, and it was enriched for nutrients responses such as nitrate and sucrose, similar to the 755 DEGs that were *HY5* dependent.

To conduct a more fine-grained analysis of gene expression patterns, we conducted hierarchical clustering and a subsequent GO enrichment analysis of the resulting clusters (Fig. 5c). Cluster 1 contained genes which increased their expression at high ambient temperature for all genotypes, and these were enriched for circadian rhythm and response to heat. This might indicate that circadian rhythm and heat sensing might be less involved downstream of *HY5*/*NRT1.1* to affect the root thermomorphogenesis and nutrient alteration at high ambient temperature. Cluster 6 contained genes that decrease their expression at high ambient temperature in Col-0 and *chl1-5,* but not in *hy5-215.* This gene set was enriched for response to nitrate and inorganic substances. Cluster 7 and 8 contained genes that decrease their expression at high ambient temperature more in *chl1-5* but less in *hy5-215*. This gene set was enriched for suberin biosynthetic process, metabolic processes such as terpenoid or organic acid, and response to water deprivation or abscisic acid. Overall, these data suggest that the *HY5-NRT1.1* regulatory mechanism controls root thermomorphogenesis and N-P alteration at high ambient temperature through the control of distinct biological processes.

### The *HY5-NRT1.1* regulatory mechanism might be conserved in soybean and rice at higher temperature

It was previously reported that HY5 and NRT1.1 homologs share similar functions in other species (Burman et al., 2018; Hu *et al*., 2019; Ji et al., 2022). Moreover, we showed that root growth responses to higher temperatures are conserved in Arabidopsis, soybean and rice (Fig. 1c, e). To explore whether the *HY5-NRT1.1* regulatory mechanism that we had discovered to be responsible for this in Arabidopsis, is conserved in soybean and rice at higher temperature, and potentially responsible for the root growth response to higher temperature, we analyzed homologs of *HY5* and *NRT1.1* in soy and rice (Fig. 6a, b). While soy has 4 homologs of both genes, *HY5* and *NRT1.1*, rice has 3 and 2 homologs of *HY5* and *NRT1.1* respectively. To explore the expression pattern of these homologs in response to high temperature, we performed qPCR (Fig. 6c, d). While all of the 4 *HY5* homologs in soybean showed a higher transcript level in the root at higher temperature, in rice, only *OsbZIP48* (*OsHY5L2*) showed increased transcript levels in the root at higher temperature. This might suggest that while all of 4 *HY5* homologs in soy have a role at higher temperature, only *OsbZIP48* might be involved in the response to higher temperature response in rice. Intriguingly, 2 *NRT1*.1 homologs in soy showed decreased transcript levels in the root at higher temperature, and the other 2 *NRT1.1* homologs in soy showed increased transcript levels in the root at higher temperature. This might suggest that the function of these genes might have diverged or neo-functionalized with regards to their role at higher temperatures. In rice, only one of 2 *NRT1.1* homologs showed a decreased transcript level in the root at higher temperature. Overall, this suggests that the role of *HY5-NRT1.1* homologs in soy is more complex, whereas *OsbZIP48-OsNRT1.1A* might be responsible for higher temperature response in rice. We performed Western blot analysis using native Arabidopsis NRT1.1 and HY5 antibody in soy and rice roots (Fig. 6e). The NRT1.1 and HY5 homolog proteins from soy and rice roots that were detected by these antibodies showed a similar pattern to Arabidopsis: NRT1.1 decreasing at higher temperatures and HY5 accumulating at higher temperature. Overall, these data suggest that *HY5-NRT1.1* regulatory mechanism might be conserved in plants and may contribute to regulate thermomorphogenesis.

**Figure 6.**
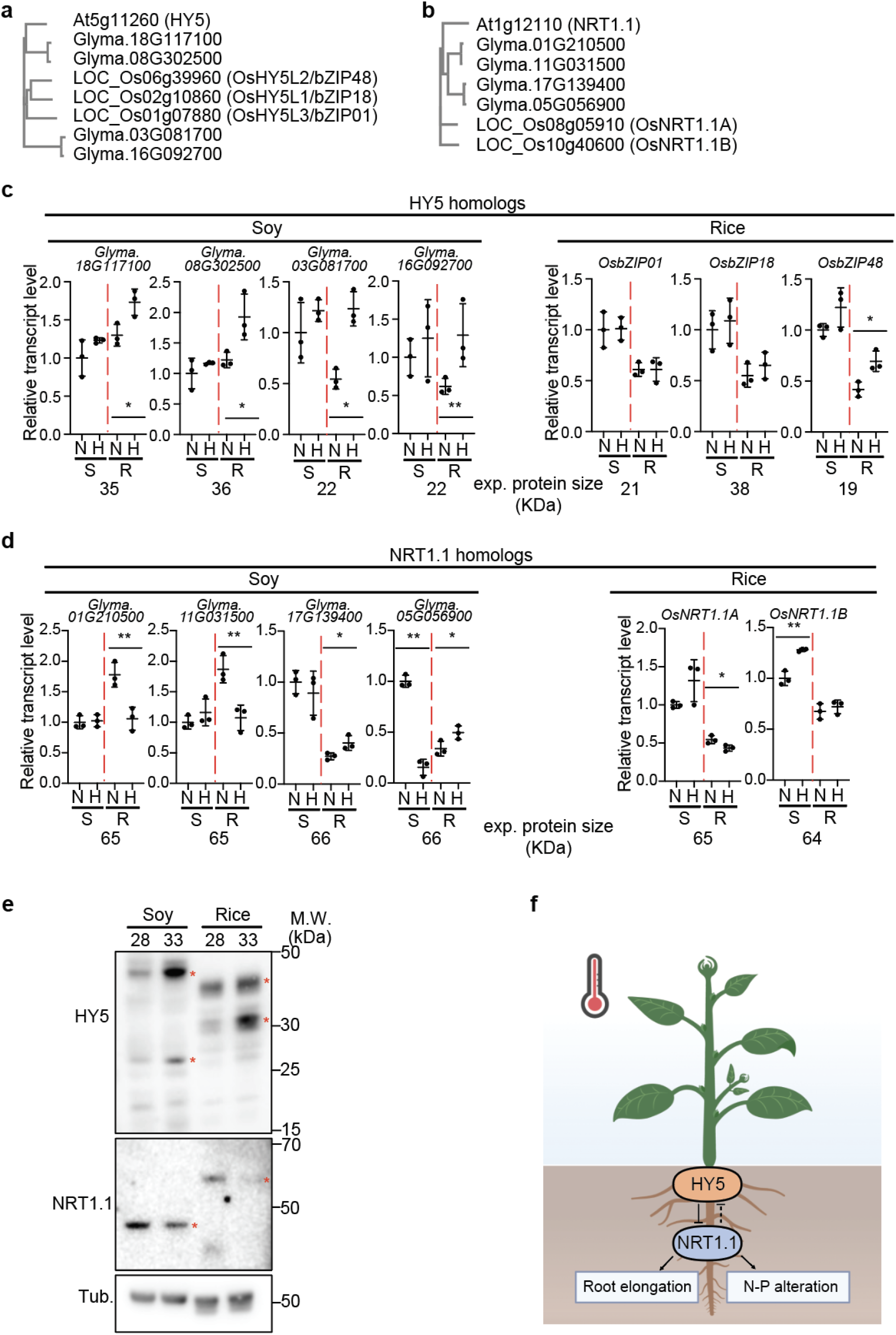
The *HY5-NRT1.1* regulatory mechanism for responses to elevated ambient temperatures is conserved in soybean and rice. **a, b** Phylogram of HY5 (**a**) and NRT1.1 (**b**) homologs in Arabidopsis, soy, and rice. **c, d** Relative transcript level of *HY5* (**c**) and *NRT1.1* (**d**) homologs in soy and rice. Relative transcript level is normalized by house-keeping genes such as rice *ubiquitin* and soy *tubulin 4*. N and H stand for Normal temperature (28°C) and Higher temperature (33°C) respectively. S and R stand for Shoot and Root respectively. Asterisks indicate statistically significant difference using one-sided Student’s t-test; *p<0.05, **p < 0.01. Expected protein size from each transcript is described below. **e** Western blot analysis using native Arabidopsis NRT1.1 and HY5 antibodies. Soybean and rice root samples were used. Red asterisks are potential HY5 and NRT1.1 bands in soybean and rice based on the expected protein size from **c** and **d**. **f** Simplified model showing root specific HY5 accumulation inhibiting NRT1.1 transcription to promote root elongation and the HY5-NRT1.1 regulatory mechanism altering N and P uptake at higher temperatures in plants. Scatter dot plots indicate mean (horizontal line) and standard deviation (error bars).

## Discussion

Thermomorphogenesis has been extensively studied in Arabidopsis. We have now shown that this response is conserved in other plant species, covering the major groups of angiosperms, monocots and dicots. We have also found a conserved interaction of thermomorphogenesis and nutrient levels. This interaction is multifaceted: thermomorphogenesis affects the nitrogen and phosphate content of plants, and it is itself dependent on sufficient levels of external nitrogen and phosphate. This interaction is governed by a *HY5*-*NRT1.1* regulatory mechanism.

The bZIP transcription factor HY5 was originally identified as a positive regulator of light signaling, and only recently emerged as a major factor to regulate temperature dependent signaling in the root. HY5 functions as both, transcription activator and repressor (Burko *et al*., 2020; Oyama *et al*., 1997; Wang *et al*., 2020; Yang et al., 2020), and regulates numerous genes by binding to its target promoter region; its binding to a distinct set of target promoters is increased at high ambient temperature (Fig. 2a-c; (Lee *et al*., 2021a). Mutation of *HY5* has wide ranging consequences for nutrient levels, not only for the levels of N and P, but also for those of Ca, Carbon, K, Na, Mn, and Zn (Supplementary Fig. 1). It will be interesting to further elucidate how HY5 changes its preferred direct targets at high ambient temperature.

Downstream of *HY5, NRT1.1* acts as a main player in the interaction of temperature responses and nutrients. NRT1.1 was originally discovered as a dual-affinity nitrate transporter and has been shown to integrate nitrate and phosphate signaling in plants (Hu *et al*., 2019; Krouk *et al*., 2010; Tsay *et al*., 1993; Ye *et al*., 2019). Our data show that decreases in both transcript and protein level of NRT1.1 are critical to trigger root responses and N-P level alterations at higher temperatures. The root response and alterations in N-P levels are conserved in Arabidopsis, soybean, and rice (Fig. 2, 3, 4, 6). It seems reasonable to assume that since NRT1.1 shares high homology among plants (Sun et al., 2014), the role of NRT1.1 might be conserved among plant species at higher temperature. Because Arabidopsis NRT1 family members have similar but diverged functions (Li et al., 2010; Lin et al., 2008; Sun *et al*., 2014), it is possible that different NRT1s or even other NRTs might have distinct roles at higher temperature. For example, *NRT2.1* has been known as a direct target of HY5 (Chen et al., 2016), however, while *NRT1.1* is strongly regulated (bound and altered transcription level) by HY5 at high ambient temperature in the root (Fig. 2a-c, supplementary Fig. 2), *NRT2.1* is not. This seems to be a similar situation as found in the PIFs. All PIFs interact with PHYTOCHROME B (PhyB) and share a redundant role in the red-light signaling pathway. However, PIF1 has a dominant role in seed germination and chlorophyll biosynthesis (Huq and Quail, 2002; Oh et al., 2006), while PIF4 and PIF7 have dominant roles in shoot thermomorphogenesis (Fiorucci *et al*., 2019; Koini *et al*., 2009), and PIF4 and PIF5 in shade avoidance response (Franklin, 2008).

Overall, the observed conserved interactions of nutrient levels and temperature responses are relevant in several ways when it comes to the ongoing climate change. Global warming will affect soil temperatures profoundly, and it has been shown that soil warming affects nutrient availability, especially N and P (Kastovska et al., 2022; Tian et al., 2023). It will therefore be important to better understand the interactions of N, P and temperature that relate to plant growth. This becomes even more important as fertilizer production and application (in particular for N) causes significant emissions of greenhouse gases (Gao and Cabrera Serrenho, 2023; Schoumans et al., 2015). While our nutrient analysis is from 4-week soil grown plants (Fig. 1g-i; Supplementary Fig. 1), longer growth and analysis of nutrients in other plant parts such as seeds might reveal additional nutrient level changes, some of which might be relevant for the nutritional value of crops. Also, effects on other nutrients at higher temperature might differ from plant species to plant species, perhaps associated with crop-specific fertilizer formulations (Hoeft, 2009; Warncke, 2009). Taken together, biotechnology or breeding strategies for taking into considerations future temperature regimes and nutrient levels will be important for overcoming the challenges that global warming will pose for the production of nutritious food and feed at the scale that is needed for a growing global population.

## Methods

### Plant materials, growth conditions, and phenotypic analyses

Arabidopsis plants were grown at LED light with light intensity of 100 μmol and diurnal condition of 16L:8D. Soy and rice plants were grown at LED light with light intensity of 200 μmol and diurnal condition of 16L:8D. Arabidopsis seedlings were plated in the media and grown vertically at either 21LJ or 28LJ for additional 5 days after 4 days of germination at 21LJ. For the media preparation, MS full media (Caisson lab, Cat. MSP33) Nitrogen deficient media (Caisson lab, Cat. MSP19), or Phosphate deficient media (Caisson lab, Cat. MSP21) were used with pH 5.7, 0.8% micropropagation Type 1 phytoagar (Caisson), and supplement of nitrogen or phosphate source according to the previous report (Gruber *et al*., 2013). And then plates were scanned through the scanner (Epson, Perfection v600) for further analysis using imageJ. Total root length of soy and rice roots was measured using Rhizovision with parameter: broken roots mode with image threshold 200 (Seethepalli et al., 2020).

### Nutrient analyses

C/N analysis and MP-AES analysis were conducted in this study. For C/N analysis, tissue samples were dried in 70LJ oven for 48 hr. Then Genogrinder (Spex SamplePrep 2010-115) was used for milling the samples. Powder samples were sent to NuMega Resonance Labs (San Diego, CA) for Perkin Elmer PE2400-Series II, CHNS/O analyzer analysis. For MP-AES analysis, Dry samples (∼5 mg) were digested with nitric acid 65% (EMD Millipore Cat. 1.00456.2500) and hydrogen peroxide 30 % (Sigma-Aldrich Cat. H3410-1L) using an Environmental Express® Hotblock digestion system (Cat. SC196). After diluting the samples with miiliQ water, they were quantified by 4210 MP-AES (Agilent). The following wavelengths were used : 202.548 nm for Zn, 214.915 nm for P, 280.271 nm for Mg, 327.395 nm for Cu, 371.993 for Fe, 403.076 nm for Mn, 589.592 nm for Na, 616.217 nm for Ca, 769.897 nm for K. The final concentration was determined using a standard curve.

### Confocal Microscopy

For the confocal microscopy experiments, plants with two genotypes, *pNRT1.1: GFP* and *pNRT1.1:NRT1.1-GFP* transgenic lines were grown at either 21LJ or 28LJ for additional 5 days after 4 days of germination at 21LJ. Zeiss LSM710 confocal microscope was used for the experiment using the 10x or 20x. Software Zeiss Zen was used for the analysis. For the PI staining, Propidium Iodide (Sigma-Aldrich, Cat. 4170) was used. Samples were gently placed on the solution and stained. For the laser and the filter, a 514 nm laser and a 520/570 nm filter for GFP. The PI signal is excited with either 488 or 514nm laser and fluorescence emission was filtered by a 600/650 nm filter.

### Protein extraction and Western blot analyses

Total protein extracts were made from 50 seedlings of root sample using 50 μL protein extraction buffer, consisting of 0.35 M Tris-Cl pH 7.5, 10x NuPAGE Sample Reducing Agent (Thermofisher, Cat. NP0009), and 4x NuPAGE LDS Sample Buffer (Thermofisher, Cat. NP0008), and 1× protease inhibitor cocktail. After boiling the samples, samples were centrifuged at 13,000 × g for 5 min and loaded into SDS-PAGE gels (Invitrogen, Cat. NP0323BOX). Separated proteins were transferred using transfer stacks (Invitrogen, Cat. IB23002) and then immunoblotted using anti-HY5 (Abiocode, Cat. R1245-2, dilution 1:3000), anti-NRT1.1 (Agrisera, Cat. AS12 2611, dilution 1:2000), or anti-Tubulin (Invitrogen, Cat. 32-2500, dilution 1:5000) antibodies. For the secondary antibodies, anti-mouse (Biorad, Cat. 170-6516, dilution 1:5000), anti-rabbit (Agrisera, Cat. AS09 602, dilution 1:5000), or were used. SuperSignal West Femto Chemiluminescent substrate (ThermoFisher Scientific, Cat. PI34094) was used for the detection of signals.

### RNA extraction, cDNA synthesis, and qRT-PCR

For RNA sample preparation, plants with three genotypes, Col-0, *hy5-215*, and *chl1-5* mutant, were grown at either 21LJ or 28LJ for additional 5 days after 4 days of germination at 21LJ. Shoot and root separated samples were collected with three independent replicates. Samples were extracted with the RNeasy Plant Mini Kit (Qiagen, Cat. 74904). Thermo Scientific™ Maxima H Minus First Strand cDNA Synthesis Kit (ThermoFisher, Cat. FERK1652) was used for cDNA synthesis. Luna® Universal qPCR Master Mix (NEB, Cat. M3003L) and qPCR machine (Biorad, CFX Opus 384 Real-Time PCR System) were used for qPCR analysis.

### RNA-seq analyses

For RNAseq sample preparation, same condition was used as qRT-PCR. The RNA quality and quantity were analyzed using a 2100 Bioanalyzer tape station (Agilent Technologies) and Qubit Fluorometer (Invitrogen). The sequencing libraries were generated by the Salk Next Generation Sequencing Core according to Illumina manufacturer’s instructions. Sequencing was performed using the Illumina Novaseq6000 platform. For RNAseq analysis, we mapped the short-reads using Arabidopsis Information resource web site (http://www.arabidopsis.org) (Berardini et al., 2015) combined with the Splice Transcripts Alignments to Reference (STAR) version 2.7.0a method (Dobin et al., 2013). Differentially Expressed Genes (DEG) analysis was performed using edgeR (Robinson et al., 2010). Critical values for the analysis are a false discovery rate (FDR < 0.05) and log2FC (> 0 or < 0). DEGs were visualized and k-means clustering method was used to classify DEGs (FDR < 0.05 and |log2FC| >1) by comparing the 21°C and 28°C treatment for Col-0 in roots via the ComplexHeatmap (Gu et al., 2016) package in R. The cluster number (k=8) was determined by sum of squared error and Bayesian information criterion. The volcano plot was created using Enhanced Volcano R package (https://github.com/kevinblighe/EnhancedVolcano).

### Chromatin Immunoprecipitation (ChIP) assays

Chromatin Immunoprecipitation (ChIP) assays were conducted as described previously with minor modifications (Lee et al., 2021b). For ChIP assay samples, 4-day-old seedlings of *pHY5:HY5-GFP* were transferred to 21 °C or 28 °C for additional 5 days and then harvested. Samples were crosslinked using 1% formaldehyde under 30 min of vacuum and 1 M glycine was added for additional 5 min for quenching. Samples were gently washed with distilled water for five times and ground thoroughly with mortar and pestle using liquid nitrogen. All the buffers for ChIP were from ChIP assay kit (Millipore, Cat. 17-295). Ground samples were placed into 1.5 mL microtube with nuclei isolation buffer for 15 min and then centrifuged at 13,000 rpm for 10 min at 4 °C. 1 mL lysis buffer was used for resuspension of the pellet. Then, sonication of chromatin pellet was performed using digital sonifier (Fisher Scientific, Sonic Dismembrator Model 500). For immunoprecipitation, ChIP grade anti-GFP (Abcam, Cat. ab6556) and dynabeads were used. After washing series of low salt wash buffer, high salt wash buffer, LiCl wash buffer, and TE buffer, we add elution buffer for elution. Samples were incubated overnight at 65 °C for reverse crosslinking after adding NaCl with final 0.2 M concentration. Finally, PCR purification kit (QIAGEN) was used for DNA purification after proteinase K treatment for 2 hours. Samples without IP were used as input DNA. Enrichment (% of input) was calculated from each sample relative to their corresponding input.

## Acknowledgements

We thank all the Busch lab members for critical discussions. We also thank Dr. Gabriel Krouk for providing *pNRT1:NRT1-GFP* seeds and helpful suggestions. The research was supported by funds from the Salk Harnessing Plants Initiative to W.B and Michigan State University to H.R.

## Author Contributions

S.L., and W.B. conceived the study and designed the experiments. S.L. and J.S. performed root phenotypic analyses. MP-AES analyses performed by G.C. RNAseq analyses performed by S.L. and L.Z. S.L. is responsible for all other experiments. All the authors analyzed the data. W.B. and H.R supervised work and provided funds and resources. S.L. and W.B. wrote the manuscript with input of all the authors. All the authors discussed the results and commented on the manuscript.

---

**Supplementary Table 1. Gene Ontology (GO) analysis of public dataset.**GO analysis was performed using public datasets including RNAseq data (Gaillochet et al., 2020, Lee et al., 2021) and ChIPseq data (Burko et al., 2020).

**Supplementary Figure 1.**
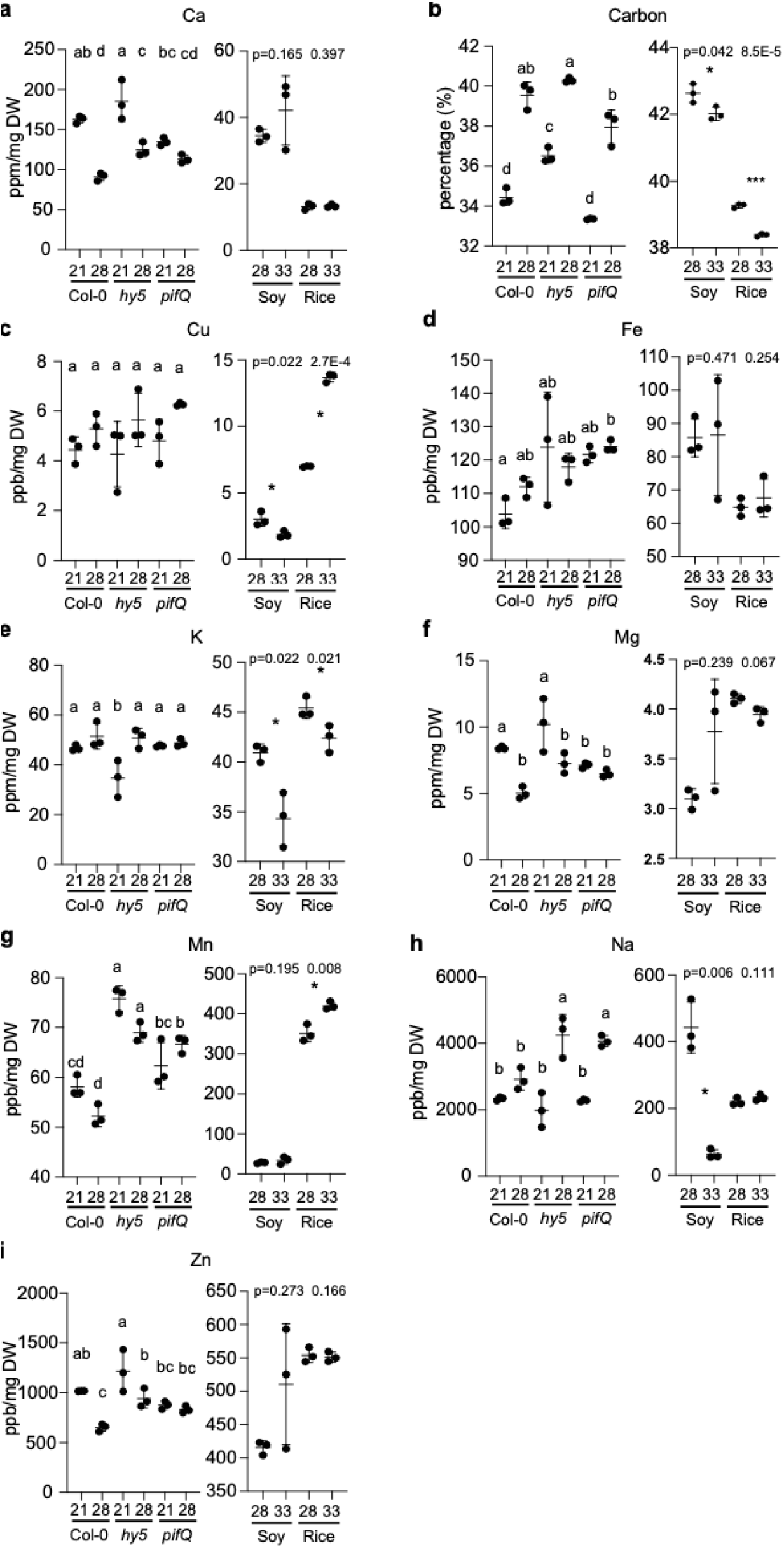
Additional nutrient analyses at higher temperatures in Arabidopsis, soybean, and rice. a-i. 9 additional nutrient analysis at higher temperature in three different species (Arabidopsis, soy, and rice) except nitrogen and phosphorus which are displayed in Fig.1 g-i. Letters a-d indicate statistically significant differences based on one-way ANOVA analysis with Tukey’s HSD test. Asterisks indicate statistically significant difference using Student’s t-test; *p<0.05, **p<0.01, and ***p<0.001. P value from one-sided Student’s t-test. Average difference of each value is indicated. Shoot parts from 4-week-old plants from Arabidopsis, soybean, and rice plants were used for the nutrient analyses. Scatter dot plots indicate mean (horizontal line) and standard deviation (error bars).

**Supplementary Figure 2.**
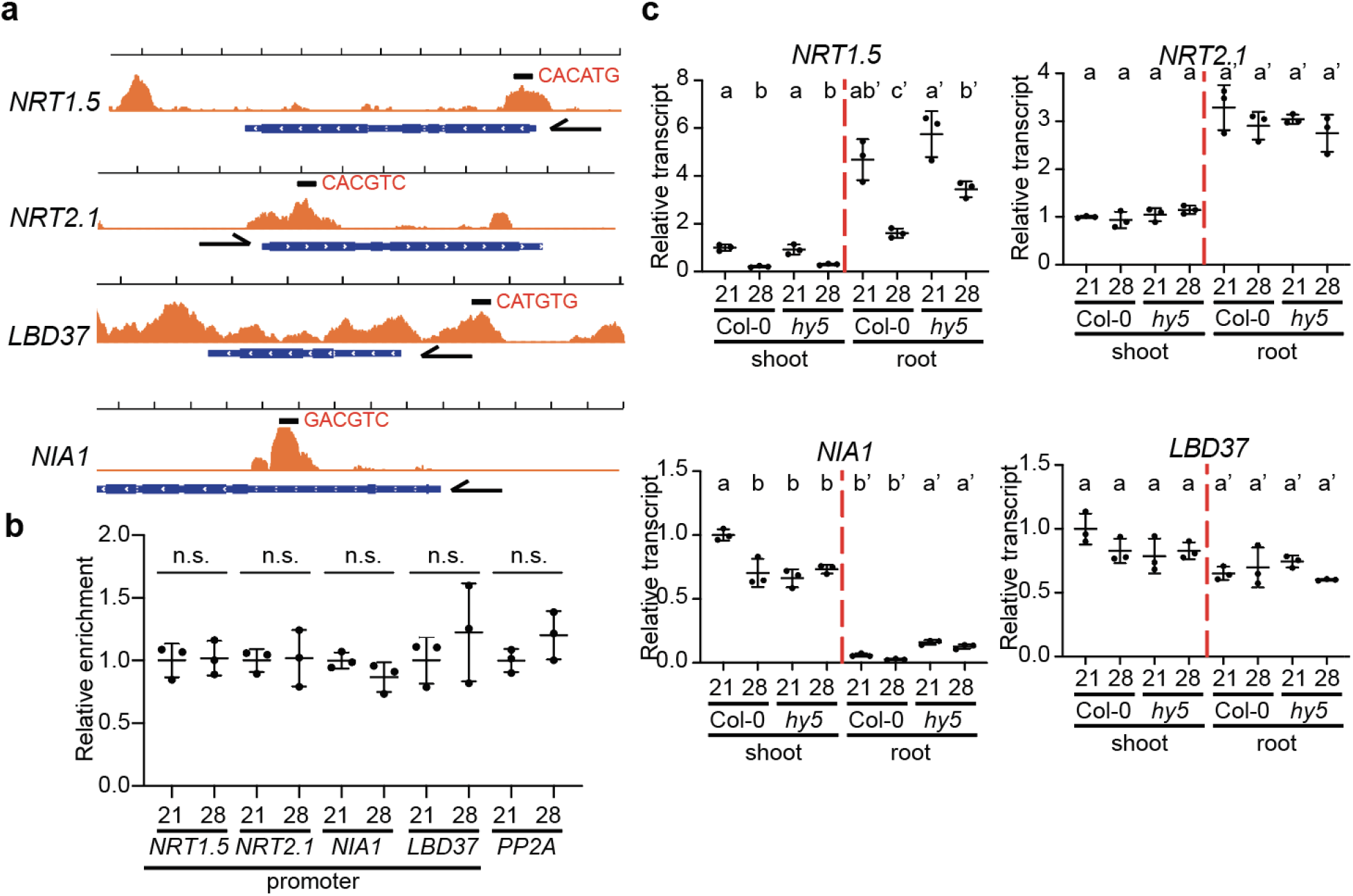
Some N-P signaling genes are not the target of HY5 at high ambient temperature. a. IGV image of HY5 ChIP-seq data from Burko et al., (2020) of N-P signaling genes with transcription direction and binding motif. **b** Scatter dot plot of ChIP-qPCR results at normal and high ambient temperature using five different genes or promoter regions. **c** Scatter dot plot of qPCR results at normal and high ambient temperature using four different genes using Col-0 and *hy5-215* seedlings with shoot and root separate samples. Relative transcript level was normalized using *PP2A* as a control and to the expression levels in the shoot. Letters a-c indicate statistically significant differences based on one-way ANOVA analysis with Tukey’s HSD test. Shoot and root samples were analyzed separately. Scatter dot plots indicate mean (horizontal line) and standard deviation (error bars)

**Supplementary Figure 3.**
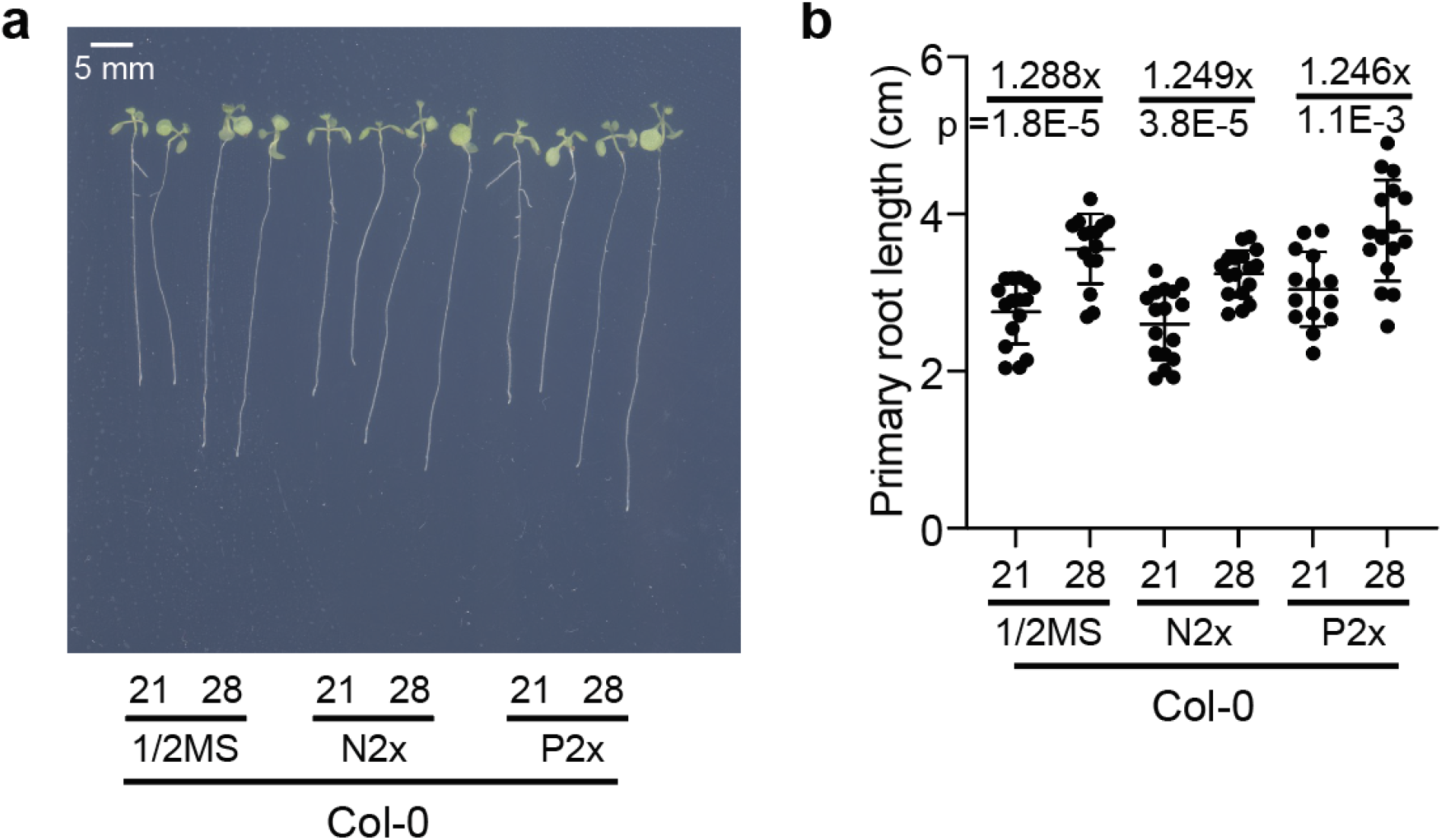
Excessive Nitrogen and Phosphorus levels do not accelerate root thermomorphogenesis. A. Phenotypes of Col-0 in different media conditions at high ambient temperature. **b** Scatter dot plot of **a**. P value from one-sided Student’s t-test. Average fold difference of each group is indicated in the top region of the plot. Scatter dot plots indicate mean (horizontal line) and standard deviation (error bars).

**Supplementary Figure 4.**
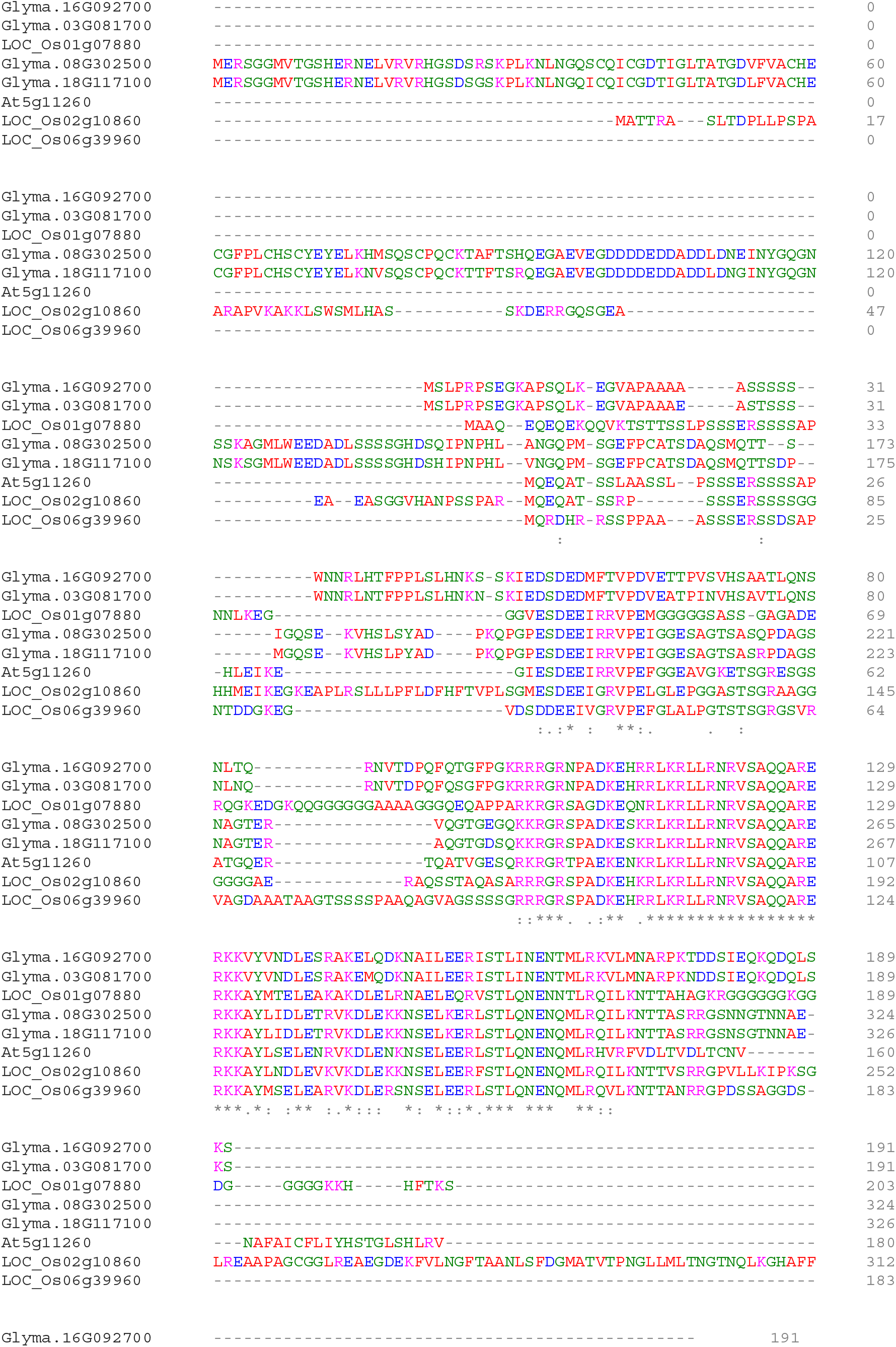
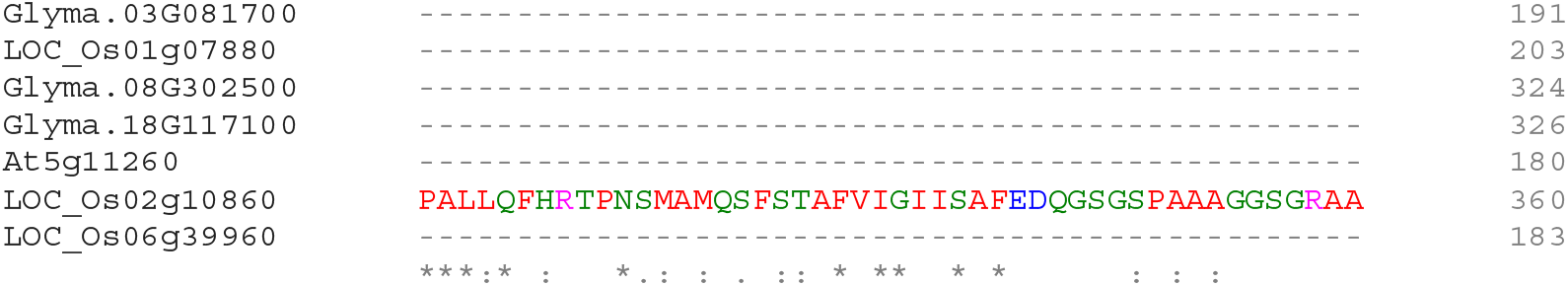
Protein alignment of HY5 homologs in Arabidopsis, soybean, and rice. Multiple protein alignment of HY5 homologs were analyzed using Clustal Omega (1.2.4). Protein sequences are from 1 in Arabidopsis (At5g11260), 4 in soybean (Glyma. 01g210500, Glyma. 11g031500, Glyma. 17g139400, and Glyma. 05g056900), and 2 in rice (LOC_Os08g05910 and LOC_Os10g40600).

**Supplementary Figure 5.**
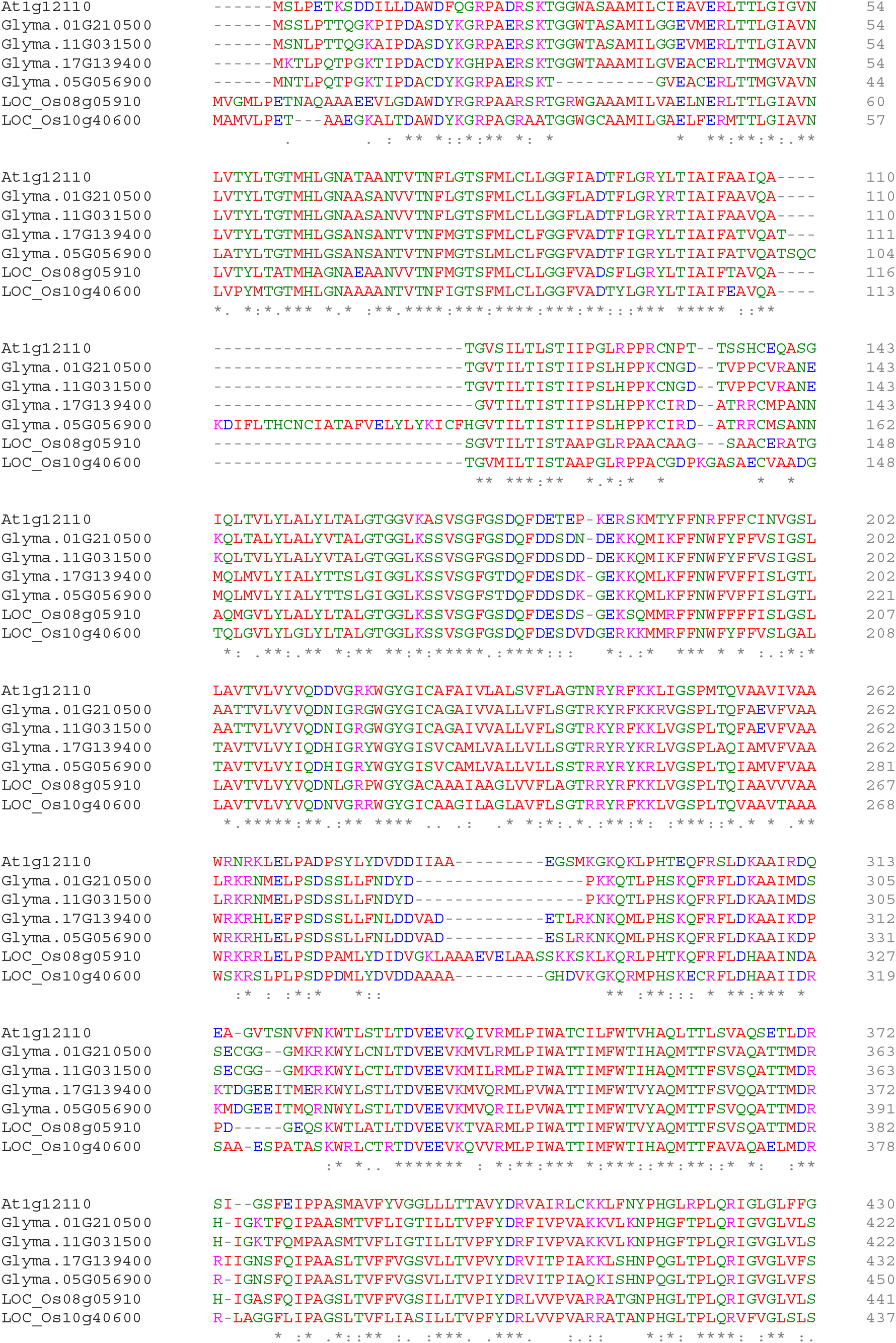
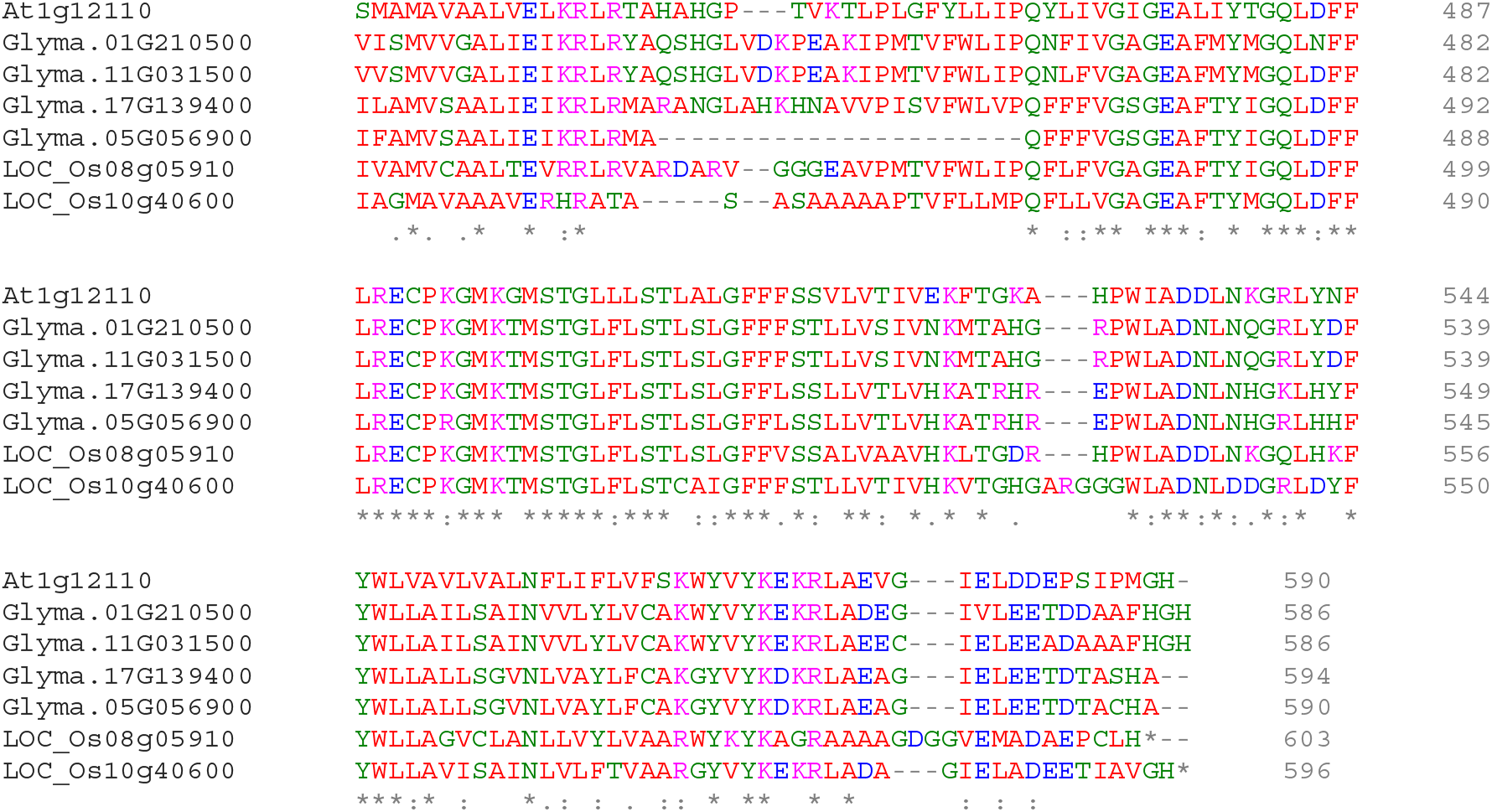
Protein alignment of NRT1.1 homologs in Arabidopsis, soybean, and rice. Multiple protein alignment of NRT1.1 homologs were analyzed using Clustal Omega (1.2.4). Protein sequences are from 1 in Arabidopsis (At1g12110), 4 in soybean (Glyma.01G210500, Glyma.11G031500, Glyma.17G139400, and Glyma.05G056900), and 2 in rice (LOC_Os08g05910 and LOC_Os10g40600).

## Notes

### Competing Interest Statement

The authors have declared no competing interest.

## References

Alvarez, J.M., Schinke, A.L., Brooks, M.D., Pasquino, A., Leonelli, L., Varala, K., Safi, A., Krouk, G., Krapp, A., and Coruzzi, G.M. (2020). Transient genome-wide interactions of the master transcription factor NLP7 initiate a rapid nitrogen-response cascade. Nat Commun 11, 1157. 10.1038/s41467-020-14979-6.

Berardini, T.Z., Reiser, L., Li, D., Mezheritsky, Y., Muller, R., Strait, E., and Huala, E. (2015). The Arabidopsis information resource: Making and mining the “gold standard” annotated reference plant genome. Genesis 53, 474–485. 10.1002/dvg.22877.

Bouain, N., Cho, H., Sandhu, J., Tuiwong, P., Prom, U.T.C., Zheng, L., Shahzad, Z., and Rouached, H. (2022). Plant growth stimulation by high CO(2) depends on phosphorus homeostasis in chloroplasts. Curr Biol 32, 4493–4500 e4494. 10.1016/j.cub.2022.08.032.

Burko, Y., Seluzicki, A., Zander, M., Pedmale, U.V., Ecker, J.R., and Chory, J. (2020). Chimeric Activators and Repressors Define HY5 Activity and Reveal a Light-Regulated Feedback Mechanism. Plant Cell 32, 967–983. 10.1105/tpc.19.00772.

Burko, Y., Willige, B.C., Seluzicki, A., Novak, O., Ljung, K., and Chory, J. (2022). PIF7 is a master regulator of thermomorphogenesis in shade. Nat Commun 13, 4942. 10.1038/s41467-022-32585-6.

Burman, N., Bhatnagar, A., and Khurana, J.P. (2018). OsbZIP48, a HY5 Transcription Factor Ortholog, Exerts Pleiotropic Effects in Light-Regulated Development. Plant Physiol 176, 1262–1285. 10.1104/pp.17.00478.

Carvalho, J.M., Barreto, R.F., Prado, R.M., Habermann, E., Branco, R.B.F., and Martinez, C.A. (2020). Elevated CO2 and warming change the nutrient status and use efficiency of Panicum maximum Jacq. PLoS One 15, e0223937. 10.1371/journal.pone.0223937.

Cassan, O., Pimpare, L.L., Dubos, C., Gojon, A., Bach, L., Lebre, S., and Martin, A. (2023). A gene regulatory network in Arabidopsis roots reveals features and regulators of the plant response to elevated CO(2). New Phytol. 10.1111/nph.18788.

Chen, C.Z., Lv, X.F., Li, J.Y., Yi, H.Y., and Gong, J.M. (2012). Arabidopsis NRT1.5 is another essential component in the regulation of nitrate reallocation and stress tolerance. Plant Physiol 159, 1582–1590. 10.1104/pp.112.199257.

Chen, X., Yao, Q., Gao, X., Jiang, C., Harberd, N.P., and Fu, X. (2016). Shoot-to-Root Mobile Transcription Factor HY5 Coordinates Plant Carbon and Nitrogen Acquisition. Curr Biol 26, 640–646. 10.1016/j.cub.2015.12.066.

Dobin, A., Davis, C.A., Schlesinger, F., Drenkow, J., Zaleski, C., Jha, S., Batut, P., Chaisson, M., and Gingeras, T.R. (2013). STAR: ultrafast universal RNA-seq aligner. Bioinformatics 29, 15–21. 10.1093/bioinformatics/bts635.

Fang, X.Z., Fang, S.Q., Ye, Z.Q., Liu, D., Zhao, K.L., and Jin, C.W. (2021). NRT1.1 Dual-Affinity Nitrate Transport/Signalling and its Roles in Plant Abiotic Stress Resistance. Front Plant Sci 12, 715694. 10.3389/fpls.2021.715694.

Fiorucci, A.S., Galvao, V.C., Ince, Y.C., Boccaccini, A., Goyal, A., Allenbach Petrolati, L., Trevisan, M., and Fankhauser, C. (2019). PHYTOCHROME INTERACTING FACTOR 7 is important for early responses to elevated temperature in Arabidopsis seedlings. New Phytol. 10.1111/nph.16316.

Franklin, K.A. (2008). Shade avoidance. New Phytol 179, 930–944. 10.1111/j.1469-8137.2008.02507.x.

G-Yull Rhee, I.J.G. (1981). The effect of environmental factors on phytoplankton growth: Temperature and the interactions of temperature with nutrient limitation. Limnology and Oceanography. doi.org/10.4319/lo.1981.26.4.0635.

Gaillochet, C., Burko, Y., Platre, M.P., Zhang, L., Simura, J., Willige, B.C., Kumar, S.V., Ljung, K., Chory, J., and Busch, W. (2020). HY5 and phytochrome activity modulate shoot-to-root coordination during thermomorphogenesis in Arabidopsis. Development 147. 10.1242/dev.192625.

Gao, Y., and Cabrera Serrenho, A. (2023). Greenhouse gas emissions from nitrogen fertilizers could be reduced by up to one-fifth of current levels by 2050 with combined interventions. Nat Food 4, 170–178. 10.1038/s43016-023-00698-w.

Gruber, B.D., Giehl, R.F., Friedel, S., and von Wiren, N. (2013). Plasticity of the Arabidopsis root system under nutrient deficiencies. Plant Physiol 163, 161–179. 10.1104/pp.113.218453.

Gu, Z., Eils, R., and Schlesner, M. (2016). Complex heatmaps reveal patterns and correlations in multidimensional genomic data. Bioinformatics 32, 2847–2849. 10.1093/bioinformatics/btw313.

Guo, F.Q., Wang, R., Chen, M., and Crawford, N.M. (2001). The Arabidopsis dual-affinity nitrate transporter gene AtNRT1.1 (CHL1) is activated and functions in nascent organ development during vegetative and reproductive growth. Plant Cell 13, 1761–1777. 10.11054/tpc.010126.

Guo, Z., Xu, J., Wang, Y., Hu, C., Shi, K., Zhou, J., Xia, X., Zhou, Y., Foyer, C.H., and Yu, J. (2021). The phyB-dependent induction of HY5 promotes iron uptake by systemically activating FER expression. EMBO Rep 22, e51944. 10.15252/embr.202051944.

Held, C., and Sadowski, G. (2016). Thermodynamics of Bioreactions. Annu Rev Chem Biomol Eng 7, 395–414. 10.1146/annurev-chembioeng-080615-034704.

Hoeft, F. (2009). Managing Soil pH and Crop Nutrients. Illinois Agronomy Handbook.

Hu, B., Jiang, Z., Wang, W., Qiu, Y., Zhang, Z., Liu, Y., Li, A., Gao, X., Liu, L., Qian, Y., et al. (2019). Nitrate-NRT1.1B-SPX4 cascade integrates nitrogen and phosphorus signalling networks in plants. Nat Plants 5, 401–413. 10.1038/s41477-019-0384-1.

Huang, L., Zhang, H., Zhang, H., Deng, X.W., and Wei, N. (2015). HY5 regulates nitrite reductase 1 (NIR1) and ammonium transporter1;2 (AMT1;2) in Arabidopsis seedlings. Plant Sci 238, 330–339. 10.1016/j.plantsci.2015.05.004.

Huq, E., and Quail, P.H. (2002). PIF4, a phytochrome-interacting bHLH factor, functions as a negative regulator of phytochrome B signaling in Arabidopsis. EMBO J 21, 2441–2450. 10.1093/emboj/21.10.2441.

Ibanez, C., Poeschl, Y., Peterson, T., Bellstadt, J., Denk, K., Gogol-Doring, A., Quint, M., and Delker, C. (2017). Ambient temperature and genotype differentially affect developmental and phenotypic plasticity in Arabidopsis thaliana. BMC Plant Biol 17, 114. 10.1186/s12870-017-1068-5.

IPCC (2022). Climate Change 2022: Impacts, Adaptation and Vulnerability. Working Group II Contribution to the IPCC Sixth Assessment Report.

Ji, H., Xiao, R., Lyu, X., Chen, J., Zhang, X., Wang, Z., Deng, Z., Wang, Y., Wang, H., Li, R., et al. (2022). Differential light-dependent regulation of soybean nodulation by papilionoid-specific HY5 homologs. Curr Biol 32, 783–795 e785. 10.1016/j.cub.2021.12.041.

Kastovska, E., Choma, M., Capek, P., Kana, J., Tahovska, K., and Kopacek, J. (2022). Soil warming during winter period enhanced soil N and P availability and leaching in alpine grasslands: A transplant study. PLoS One 17, e0272143. 10.1371/journal.pone.0272143.

Koini, M.A., Alvey, L., Allen, T., Tilley, C.A., Harberd, N.P., Whitelam, G.C., and Franklin, K.A. (2009). High temperature-mediated adaptations in plant architecture require the bHLH transcription factor PIF4. Curr Biol 19, 408–413. 10.1016/j.cub.2009.01.046.

Krouk, G., Lacombe, B., Bielach, A., Perrine-Walker, F., Malinska, K., Mounier, E., Hoyerova, K., Tillard, P., Leon, S., Ljung, K., et al. (2010). Nitrate-regulated auxin transport by NRT1.1 defines a mechanism for nutrient sensing in plants. Dev Cell 18, 927–937. 10.1016/j.devcel.2010.05.008.

Kumar, S.V., Lucyshyn, D., Jaeger, K.E., Alos, E., Alvey, E., Harberd, N.P., and Wigge, P.A. (2012). Transcription factor PIF4 controls the thermosensory activation of flowering. Nature 484, 242–245. 10.1038/nature10928.

Lau, O.S., Song, Z., Zhou, Z., Davies, K.A., Chang, J., Yang, X., Wang, S., Lucyshyn, D., Tay, I.H.Z., Wigge, P.A., and Bergmann, D.C. (2018). Direct Control of SPEECHLESS by PIF4 in the High-Temperature Response of Stomatal Development. Curr Biol 28, 1273–1280 e1273. 10.1016/j.cub.2018.02.054.

Lee, J., He, K., Stolc, V., Lee, H., Figueroa, P., Gao, Y., Tongprasit, W., Zhao, H., Lee, I., and Deng, X.W. (2007). Analysis of transcription factor HY5 genomic binding sites revealed its hierarchical role in light regulation of development. Plant Cell 19, 731–749. 10.1105/tpc.106.047688.

Lee, S., Wang, W., and Huq, E. (2021a). Spatial regulation of thermomorphogenesis by HY5 and PIF4 in Arabidopsis. Nat Commun 12, 3656. 10.1038/s41467-021-24018-7.

Lee, S., Zhu, L., and Huq, E. (2021b). An autoregulatory negative feedback loop controls thermomorphogenesis in Arabidopsis. PLoS Genet 17, e1009595. 10.1371/journal.pgen.1009595.

Li, J.Y., Fu, Y.L., Pike, S.M., Bao, J., Tian, W., Zhang, Y., Chen, C.Z., Zhang, Y., Li, H.M., Huang, J., et al. (2010). The Arabidopsis nitrate transporter NRT1.8 functions in nitrate removal from the xylem sap and mediates cadmium tolerance. Plant Cell 22, 1633–1646. 10.1105/tpc.110.075242.

Lin, S.H., Kuo, H.F., Canivenc, G., Lin, C.S., Lepetit, M., Hsu, P.K., Tillard, P., Lin, H.L., Wang, Y.Y., Tsai, C.B., et al. (2008). Mutation of the Arabidopsis NRT1.5 nitrate transporter causes defective root-to-shoot nitrate transport. Plant Cell 20, 2514–2528. 10.1105/tpc.108.060244.

Martins, S., Montiel-Jorda, A., Cayrel, A., Huguet, S., Roux, C.P., Ljung, K., and Vert, G. (2017). Brassinosteroid signaling-dependent root responses to prolonged elevated ambient temperature. Nat Commun 8, 309. 10.1038/s41467-017-00355-4.

Medici, A., Marshall-Colon, A., Ronzier, E., Szponarski, W., Wang, R., Gojon, A., Crawford, N.M., Ruffel, S., Coruzzi, G.M., and Krouk, G. (2015). AtNIGT1/HRS1 integrates nitrate and phosphate signals at the Arabidopsis root tip. Nat Commun 6, 6274. 10.1038/ncomms7274.

Oh, E., Yamaguchi, S., Kamiya, Y., Bae, G., Chung, W.I., and Choi, G. (2006). Light activates the degradation of PIL5 protein to promote seed germination through gibberellin in Arabidopsis. Plant J 47, 124–139. 10.1111/j.1365-313X.2006.02773.x.

Oyama, T., Shimura, Y., and Okada, K. (1997). The Arabidopsis HY5 gene encodes a bZIP protein that regulates stimulus-induced development of root and hypocotyl. Genes Dev 11, 2983–2995. 10.1101/gad.11.22.2983.

Remans, T., Nacry, P., Pervent, M., Girin, T., Tillard, P., Lepetit, M., and Gojon, A. (2006). A central role for the nitrate transporter NRT2.1 in the integrated morphological and physiological responses of the root system to nitrogen limitation in Arabidopsis. Plant Physiol 140, 909–921. 10.1104/pp.105.075721.

Robinson, M.D., McCarthy, D.J., and Smyth, G.K. (2010). edgeR: a Bioconductor package for differential expression analysis of digital gene expression data. Bioinformatics 26, 139–140. 10.1093/bioinformatics/btp616.

Schleuning, M., Frund, J., Schweiger, O., Welk, E., Albrecht, J., Albrecht, M., Beil, M., Benadi, G., Bluthgen, N., Bruelheide, H., et al. (2016). Ecological networks are more sensitive to plant than to animal extinction under climate change. Nat Commun 7, 13965. 10.1038/ncomms13965.

Schoumans, O.F., Bouraoui, F., Kabbe, C., Oenema, O., and van Dijk, K.C. (2015). Phosphorus management in Europe in a changing world. Ambio 44 Suppl 2, S180–192. 10.1007/s13280-014-0613-9.

Seethepalli, A., Guo, H., Liu, X., Griffiths, M., Almtarfi, H., Li, Z., Liu, S., Zare, A., Fritschi, F.B., Blancaflor, E.B., et al. (2020). RhizoVision Crown: An Integrated Hardware and Software Platform for Root Crown Phenotyping. Plant Phenomics 2020, 3074916. 10.34133/2020/3074916.

Sun, J., Bankston, J.R., Payandeh, J., Hinds, T.R., Zagotta, W.N., and Zheng, N. (2014). Crystal structure of the plant dual-affinity nitrate transporter NRT1.1. Nature 507, 73–77. 10.1038/nature13074.

Tian, Y., Shi, C., Malo, C.U., Kwatcho Kengdo, S., Heinzle, J., Inselsbacher, E., Ottner, F., Borken, W., Michel, K., Schindlbacher, A., and Wanek, W. (2023). Long-term soil warming decreases microbial phosphorus utilization by increasing abiotic phosphorus sorption and phosphorus losses. Nat Commun 14, 864. 10.1038/s41467-023-36527-8.

Tsay, Y.F., Schroeder, J.I., Feldmann, K.A., and Crawford, N.M. (1993). The herbicide sensitivity gene CHL1 of Arabidopsis encodes a nitrate-inducible nitrate transporter. Cell 72, 705–713. 10.1016/0092-8674(93)90399-b.

Vu, L.D., Xu, X., Gevaert, K., and De Smet, I. (2019). Developmental Plasticity at High Temperature. Plant Physiol 181, 399–411. 10.1104/pp.19.00652.

Wang, X., Wang, H.F., Chen, Y., Sun, M.M., Wang, Y., and Chen, Y.F. (2020). The Transcription Factor NIGT1.2 Modulates Both Phosphate Uptake and Nitrate Influx during Phosphate Starvation in Arabidopsis and Maize. Plant Cell 32, 3519–3534. 10.1105/tpc.20.00361.

Warncke, D., J. Dahl, and L. Jacobs (2009). Nutrient Recommendations for field crops in Michigan. Bull. E2904. . Michigan State Univ. Ext., East Lansing, MI.

Yang, C., Shen, W., Yang, L., Sun, Y., Li, X., Lai, M., Wei, J., Wang, C., Xu, Y., Li, F., et al. (2020). HY5-HDA9 Module Transcriptionally Regulates Plant Autophagy in Response to Light-to-Dark Conversion and Nitrogen Starvation. Mol Plant 13, 515–531. 10.1016/j.molp.2020.02.011.

Ye, J.Y., Tian, W.H., and Jin, C.W. (2019). A reevaluation of the contribution of NRT1.1 to nitrate uptake in Arabidopsis under low-nitrate supply. FEBS Lett 593, 2051–2059. 10.1002/1873-3468.13473.

Yu, X., Sukumaran, S., and Mrton, L. (1998). Differential expression of the arabidopsis nia1 and nia2 genes. cytokinin-induced nitrate reductase activity is correlated with increased nia1 transcription and mrna levels. Plant Physiol 116, 1091–1096. 10.1104/pp.116.3.1091.

